# Stable Individual Differences from Dynamic Patterns of Function: Brain Network Flexibility Predicts Openness/Intellect and Intelligence

**DOI:** 10.1101/2024.01.05.574386

**Authors:** Tyler A. Sassenberg, Adam Safron, Colin G. DeYoung

## Abstract

Growing understanding of the nature of brain function has led to increased interest in interpreting the properties of large-scale brain networks. Methodological advances in network neuroscience provide means to decompose these networks into smaller functional communities and measure how they reconfigure over time as an index of their dynamic and flexible properties. Recent evidence has identified associations between flexibility and a variety of traits pertaining to complex cognition including creativity and working memory. The present study used measures of dynamic resting-state functional connectivity in data from the Human Connectome Project (*N* = 994) to test associations with Openness/Intellect and general intelligence, two traits that involve flexible cognition. Using a machine-learning cross-validation approach, we identified reliable associations of intelligence with cohesive flexibility of parcels in large communities across the cortex, and of Openness/Intellect with overall flexibility among parcels in smaller communities. These findings are reasonably consistent with previous theories of the neural correlates of intelligence and Openness/Intellect, and help to expand on previous associations of behavior and dynamic functional connectivity within the context of broader personality dimensions.

## Stable Individual Differences from Dynamic Patterns of Function: Brain Network Flexibility Predicts Openness/Intellect and Intelligence

Neuroscience research using techniques like functional magnetic resonance imaging (fMRI) to understand the structure and function of the brain has led to a growing consensus regarding the existence of broad macroscale brain networks. Instead of conceptualizing brain functions as localized in discrete regions, a network-based paradigm has emerged from the observation that blood-oxygen-level dependent (BOLD) signals across the brain tend to exhibit widespread reliable correlations that are notably similar in resting-state and task conditions (Eickhoff et al. 2011; Yeo et al. 2011; Smith et al. 2013). These functional connections between spatially distant regions allow identification of a canonical set of large-scale functional networks, each with unique behavioral correlates.

To capture the complexity of interactions within and between these brain networks, measures of functional connectivity are often derived using multivariate approaches like independent components analysis, which identifies patterns of intercorrelations among voxel timeseries and maps them to statistically independent components in the brain (Hyvӓrinen and Oja 2000). Another common approach is the use of graph theory metrics to characterize regions of interest as nodes and their correlations as edges, which together can provide a variety of measures of network topology (Bullmore and Sporns 2012; Rubinov and Sporns 2010). One particularly useful graph theory metric is community structure, which describes the organization of broad networks into smaller functional units called communities (also called cliques or modules), that tend to exhibit greater connectivity within their unit than with the rest of the network (Porter et al. 2009).

Much of the research seeking to identify patterns in brain organization has relied on timeseries across the duration of an entire fMRI scan to produce estimates of typical connectivity patterns in the brain, described as *static* functional connectivity. However, a growing emphasis on understanding the brain as a complex dynamical system has emphasized a form of connectivity reflecting variation in temporal dynamics, or *dynamic* functional connectivity. A variety of methods have been developed to estimate dynamic functional connectivity, including characterizing the most predictive aspects of BOLD timeseries (Karahanoğlu and Van De Ville 2015; Tagliazucchi et al. 2012), constraining functional pathways by structural connections (Griffa et al. 2017; Vohryzek et al. 2020), describing connectivity through metastates or trajectories through attractor landscapes (Breakspear 2017; Vidaurre et al. 2017), and using sliding-window approaches (Allen et al. 2014; Preti et al. 2017). An important advantage of dynamic functional connectivity analysis is the ability to capture the evolution of cortical topology measures like community structure over time as a potential index of moment-to-moment brain state transitions.

The variety of approaches for studying dynamic functional connectivity offers a framework for understanding the neural correlates of personality traits. Recent research has speculated that indices of network flexibility, or the tendency of brain regions to change from one community to another, may be reliable measures of adaptive functioning, and may demonstrate associations with the personality metatrait Plasticity (Safron et al. 2022). Plasticity reflects the shared variance of the traits Extraversion and Openness/Intellect, and describes individual differences in exploration, adaptability, and the tendency to develop new goals, strategies, and interpretations (DeYoung 2006, 2015). Safron et al. (2022) speculated that the patterns of cognition and behavior described by Plasticity may be a consequence of particular kinds of network flexibility. Specifically, they hypothesized that cohesive flexibility, describing the tendency for functional regions to change communities together, may be a good index of self-organizing criticality within the brain and may be positively associated with the general exploratory tendencies described by Plasticity.

However, much of the evidence used to support this hypothesis involves associations between network flexibility and a variety of lower-order facets specifically within the Openness/Intellect domain. Openness/Intellect is the Big Five dimension that describes individual differences in cognitive exploration, or variation in the tendency to seek out, detect, appreciate, understand, and use sensory and abstract information (DeYoung 2015). Many features of complex cognition have been shown to be associated with network flexibility, including working memory capacity (Braun et al. 2015), cognitive fatigue (Betzel, Satterthwaite et al. 2017), need for cognition (an excellent marker of the Intellect aspect of Openness/Intellect), and creativity (He et al. 2019; Patil et al. 2021). These findings suggest the possibility that network flexibility may be more simply related to traits occupying positions in the personality hierarchy lower than Plasticity, such as Openness/Intellect or intelligence. This notion is also supported by evidence describing associations of facets of Openness/Intellect like creativity and intelligence with processing speed and the capacity for organizing brain functional dynamics across broad networks (Lee and Chabris 2013; Langer et al. 2011; Zhuang et al. 2021). Finally, if the association were with Plasticity, network flexibility should be similarly related to Extraversion as it is to Openness/Intellect, and this seems unlikely because Extraversion has been most linked to reward sensitivity and more specific neural processes in reward networks that are not as broadly distributed throughout the cortex as networks supporting complex cognition (Wacker and Smillie 2015; Smillie et al. 2019).

In the present research, therefore, we draw inspiration from the hypothesis of Safron et al. (2022) to test associations between flexibility in patterns of dynamic resting-state functional connectivity with personality, but we hypothesize that associations are more likely with Openness/Intellect and general intelligence than with Plasticity more generally. Using a machine learning approach with publicly available resting-state data from the Human Connectome Project, we perform what is, to the best of our knowledge, the first empirical test of associations of brain network flexibility with two influential constructs that occupy different strata of the personality hierarchy but share outcomes pertaining to engagement with new and complex information. (Because of our doubts about the relevance of Plasticity as a criterion personality trait, and because our analytic approach did not allow modeling Plasticity as a latent variable (which is the preferred method for measuring the metatraits), we focus our analyses on Openness/Intellect and intelligence. Analyses using metatrait Plasticity as a manifest variable are reported in the online supplement and did not show any significant effects.)

## Methods

### Participants

A total of 994 participants (528 female) were selected from the WU-Minn Consortium of the Human Connectome Project (HCP) (Van Essen et al. 2012). Initial HCP exclusion criteria included a history of severe psychiatric, neurological, or medical disorders. Participants of the 1200 young adult sample were further excluded from subsequent analyses on the basis of missing personality and intelligence task data, and missing resting-state fMRI scan data.

Participants’ ages ranged from 22 to 37 years old (*M* = 28.7, *SD* = 3.7). All participants of the 1200 young adult sample provided informed consent. Further information regarding the informed consent procedure is described by Van Essen et al. (2013). All study protocols were approved by the Institutional Review Board of Washington University in St. Louis. Data are available from the HCP website: https://db.humanconnectome.org

### Behavioral Measures

#### Big Five

Participants included in the present study completed the NEO Five-Factor Inventory (FFI). The NEO-FFI is a short form of the NEO Personality Inventory, Revised (NEO PI-R; Costa and McCrae 1992), consisting of 12 items per factor. Scale scores were calculated as item averages, using a five-point Likert scale ranging from 0 (*strongly disagree*) to 4 (*strongly agree*). Openness/Intellect scores were used in subsequent analyses, and scores of the remaining Big Five were included as covariates.

#### Intelligence

In line with previous research assessing brain-behavior associations in HCP, a measure of general intelligence (*g*) was calculated by entering 10 cognitive tasks from the NIH Toolbox (Heaton et al. 2014) and Penn Computerized Neurocognitive Battery (Moore et al. 2015) into an exploratory bifactor model (Dubois et al. 2018a; Feilong et al. 2021). These tasks included the Oral Reading Recognition (*ReadEng_Unadj*), Picture Vocabulary (*PicVocab_Unadj*), Picture Sequence Memory (*PicSeq_Unadj*), Flanker Inhibitory Control and Attention (*Flanker_Unadj*), Dimensional Change Card Sort (*CardSort_Unadj*), Pattern Comparison Processing Speed (*ProcSpeed_Unadj*), Penn Progressive Matrices (*PMAT24_A_CR*), Penn Word Memory Test (*IWRD_TOT*), Variable Short Penn Line Orientation Test (*VSPLOT_TC*), and List Sorting Tasks (*ListSort_Unadj*). Consistent with previous literature, parallel analysis suggested a structure with four group factors (Dubois et al. 2018a), and this approach produced a well-fitting model with a general factor (*g*) explaining 56% percent of the variance among task scores (RMSEA = .03, CFI = .98, SRMR = .02). Factor scores for *g* were computed to be used in subsequent analyses.

### fMRI Data Acquisition and Preprocessing

fMRI data were acquired using a customized 3T Siemens Skyra scanner for all participants at Washington University in St. Louis. The present study used a single left-to-right phase encoded resting-state scan acquired using the following parameters: 72 axial slices; TR = 0.4 s; TE = 33 ms; flip angle = 52°; multiband acceleration factor = 8; voxel dimensions = 2 x 2 x 2 mm^3^; pixel bandwidth = 2,290 Hz. Additionally, high-resolution T1-weigthed MPRAGE structural images were acquired for anatomical surface registration with the following parameters: TR = 24 s; TE = 2.14 ms; flip angle = 8°; voxel dimensions = 0.7 x 0.7 x 0.7 mm^3^. Resting-state scans were preprocessed using the HCP minimal preprocessing pipeline and motion artifacts were removed using ICA-FIX (Burgess et al. 2016). Relative mean framewise displacement was also computed to be included as a covariate in subsequent analyses. Details of the HCP minimal preprocessing pipeline are described in greater depth in previous literature (Glasser et al. 2013; Ugurbil et al. 2013).

### Group Prior Individualized Parcellation

Functional regions were identified using an individualized cortical parcellation approach.

For each participant, ICA-denoised resting-state fMRI scans in subject-native space were first resampled to a common cortical surface mesh, and the BOLD timeseries at each vertex were normalized to zero mean and unit variance. The resulting subject surface data were then overlaid with a pre-defined group atlas with 400 functionally distinct regions (Schaefer et al. 2018) mapped to the 17-network atlas defined by Yeo et al. (2011). A Bayesian algorithm was applied to iteratively adjust parcel boundaries to maximize within-parcel homogeneity according to each participant’s unique patterns of functional connectivity (Chong et al. 2017). This algorithm was applied across 20 iterations so that participants had no more than one vertex on the cortical surface mesh changing its parcel identification on the final iteration, ensuring stable modifications of parcel boundaries. Through this process, each participant acquired a unique variant of a standard group-level atlas, such that the boundaries of the parcels of the initial atlas optimally reflected each individual’s patterns of resting-state functional connectivity, while parcels maintained their identity across subjects. Code used to produce individualized parcels of the Schaefer atlas is available at https://neuroimageusc.github.io/GPIP

### Dynamic Network Construction

Dynamic functional connectivity matrices were identified using a sliding window approach. For each participant, the average timeseries of each of their individualized parcels were divided into a set of 16 windows of 75 TRs (approximately 54 seconds). This division was motivated by previous research describing the susceptibility of windows shorter than 40 seconds to sampling variability and obscuring genuine patterns of network reconfiguration (Zalesky and Breakspear 2015; Leonardi and Van De Ville 2015). Additionally, this window size permitted windows of equal length across the duration of the scan. To assess functional connectivity between each pair of parcels, we constructed a low frequency wavelet coherence matrix describing the magnitude squared coherence of the scale two Daubechies wavelet (length 4) decomposition of the timeseries for each pair of parcels within the band-pass filtered range of .125-.25 Hz in each window using the WMTSA Wavelet Toolkit in MATLAB (https://www.atmos.washington.edu/wmtsa/). Magnitude squared coherence is a measure of functional connectivity that describes similarity in the signal waveform within a consistent frequency range. Using this metric ensures a stable index of functional connectivity across windows, while also conferring greater robustness to noise and signal outliers compared to product-moment correlation measures of dynamic functional connectivity (Andrew and Pfurtscheller, 1996; Zhang et al. 2016). Additionally, previous research has demonstrated that low frequency wavelet analyses are well-suited for measuring the properties of signals that typify resting-state activity in the cortex using fMRI (Maxim et al. 2005). These coherence matrices were concatenated across windows to create a single 400 × 400 × 16 array for each participant.

#### Community Identification

To quantify the temporal reconfiguration of communities of parcels, we utilized a greedy Louvain multi-layer modularity algorithm to assign all parcels in all layers to communities, where each participant’s windowed 400 × 400 coherence matrix was used as a single layer in a multi-layer network (Jutla et al. 2011; Mucha et al. 2010; Betzel et al. 2019). Multi-layer modularity is described by the maximization of the modularity quality function:

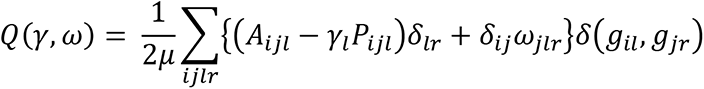

where *Aijl* describes the coherence between nodes *i* and *j* in layer *l*. The quantity *δ*(*g_il_*, *g_jr_*), known as the Kronecker delta, describes the community assignment of nodes across layers.

This algorithm defines communities as clusters of nodes characterized by stronger connections than would be expected by chance, and describes these communities by the density of these connections (Girvan and Newman 2002; Newman 2012). The function is modulated by two tuning parameters: the structural resolution of the communities, γ, and the temporal resolution of the communities, ω. Maximizing the function for small values of γ results in the identification of a few large communities, and maximizing the function for large values of γ results in the identification of many small communities. Similarly, maximizing the function for small values of ω results in greater variability in community structure over time, and large values of ω permit greater community homogeneity over time.

The use of the multi-layer modularity algorithm also relies on the specification of a suitable null model, *Pijl*, with which to compare the configuration of observed connections. In the present analyses, we utilized the multi-layer modification of the common Newman-Girvan model (Porter et al. 2009). This model was chosen on account of its ability to test the hypothesis that the observed network’s communities are a function of the particular degree sequence, and for its utility when testing networks in which connections can occur between any pair of nodes (Betzel, Medaglia et al. 2017). (Code for identifying communities using multi-layer modularity maximization is available at https://www.brainnetworkslab.com/coderesources.)

We specified γ = 1 and ω = 1 in the present analyses, in line with previous research (Bassett et al. 2013; Betzel, Satterthwaite et al. 2017). While there are more and less common approaches to setting these tuning parameters, there is no singular canonical method, and different parameter settings may be appropriate for different datasets, and potentially for capturing differing dynamics within a single experiment. Additionally, because we did not know how many communities would be identified using γ = 1, and to investigate associations of Openness/Intellect and intelligence with functional communities of different sizes, we also specified partitions using γ = 1.1 and 1.2. These partitions produced functionally recognizable communities, and exhibited sufficient variability across windows upon visual inspection. An example of each of these three community partitions mapped to the atlas by Schaefer et al. (2018) in one subject is illustrated in Figure 1. These patterns were similar across subjects.

**Figure 1.**
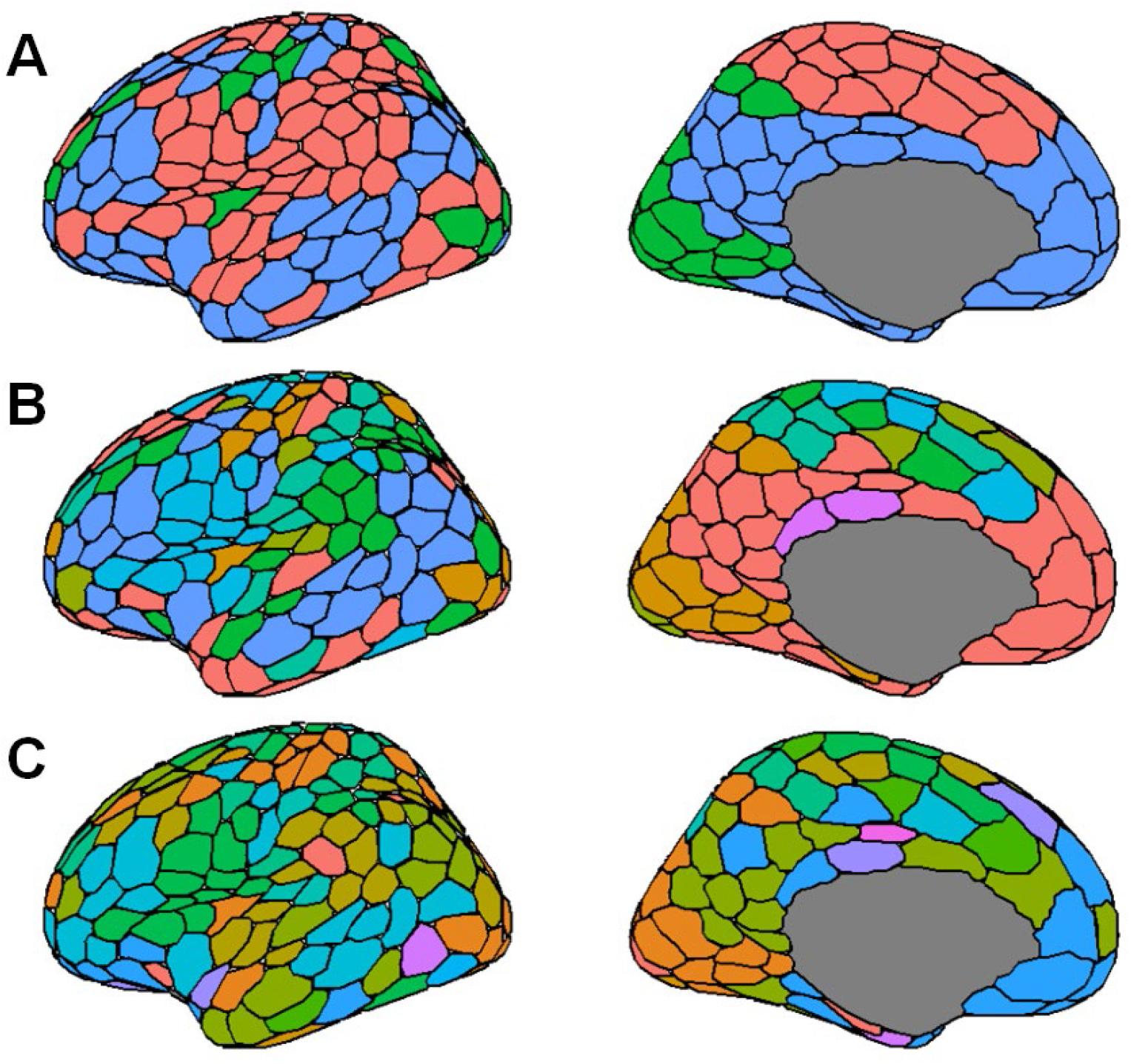
Three community structural partitions in subject 100206 at the first time window. Colors correspond to unique functional communities identified through multi-layer modularity maximization at γ = 1 (**A**), 1.1 (**B**), and 1.2 (**C**). Colors signify unique communities only within each partition, and do not represent equivalent communities across panels.

### Network Flexibility

For each participant, we computed measures of overall node flexibility, cohesive flexibility, and disjoint flexibility using the Network Community Toolbox in MATLAB (http://commdetect.weebly.com). Overall flexibility is defined as the proportion of the number of times a node changes communities to the number of times a node could change communities.

Cohesive flexibility is defined as the proportion of the number of times a node moves *with other nodes* from one community to another, relative to the number of times it could change communities. Disjoint flexibility is defined as the proportion of the number of times a node changes communities *by itself* to the number of possible times it could change communities (Bassett et al. 2011; Telesfold et al. 2017). Since this multi-layer modularity optimization algorithm relies on randomized components, we repeated the community identification procedure 10 times for each participant and averaged the values for each type of flexibility measure across all iterations to produce a final value for each parcel.

### Analysis

#### Predicting Openness/Intellect from Parcel Flexibility

We used elastic net regression to identify potential associations of Openness/Intellect with flexibility metrics of potentially influential parcels. To assess the predictive ability of the flexibility of these parcels, the original sample was divided into equally sized training and test samples (*N*’s = 497). Where multiple individuals from a single family were included in the sample, all individuals within a family were assigned to the same subsample, ensuring complete independence of the training and test samples.

Flexibility measures for each of the 400 individualized parcels were included as predictors in the model. Age, gender, mean relative RMS movement, Conscientiousness, Agreeableness, Neuroticism, Extraversion, and *g* were included as covariates. We also regressed out the effects of handedness, total brain volume (*FS_BrainSeg_Vol*), and the type of image reconstruction algorithm used in the HCP minimal preprocessing pipeline, which changed during HCP data collection (*fMRI_3T_ReconVrs*). These covariates have previously been demonstrated to affect global functional connectivity patterns in resting-state fMRI (Pool et al. 2015; Li et al. 2015; Van Dijk et al. 2012; Elam 2015). All covariates were regressed out of flexibility measures and Openness/Intellect using multiple linear regression.

Elastic net models were fit using the *glmnet* package in R (Friedman et al. 2010). Flexibility measures were log-transformed and standardized before entering the model. The mixing (α) and penalty parameters (λ) were optimized in the training sample with 10-fold cross validation for each model through a grid search using the *caret* package in R (Kuhn 2021). The model with parameters producing the highest *R*^2^ value across folds was selected for further analyses. Coefficients derived in the training sample were then applied to the predictors in the test sample. This procedure was then repeated using measures of cohesive and disjoint flexibility, across each community structural partition. Measures of root-mean squared error (RMSE), mean absolute error (MAE), and *R*^2^ were calculated for both samples to assess model fit. In line with previous research, the *R*^2^ values in the test sample were computed as the squared correlation between the observed scores and fitted values. Nonparametric permutation testing was used to determine the significance of model performance in the test set. A null distribution of *R*^2^ values was created by shuffling the flexibility measures in the test set to break any dependence between the personality scores and flexibility measures. This was accomplished using Freedman-Lane permutation to properly account for the role of covariates (Freedman and Lane 1983), and with exchangeability blocks to account for family structure within the test set (Winkler et al. 2014). This null distribution was then used to determine whether the variance explained in the test set was significantly greater than expected by chance.

#### Predicting Intelligence from Parcel Flexibility

The same procedure was used to predict *g*. These models used age, gender, mean relative RMS movement, handedness, total brain volume, and the type of image reconstruction algorithm as covariates. Separate models were fit for each of the three types of flexibility across all community partitions.

#### Analytic Procedure and Influence of Covariates

To address the influence of particular analytic choices on the predictive ability of these models, models predicting Openness/Intellect were fit without the remaining Big Five, and again without the effect of intelligence to determine how much the choice of covariates had on the hypothesized results. Similarly, models predicting intelligence were also fit including the Big Five as covariates. To further test the specificity of the prediction, we also repeated these analyses using each of the Big Five as criterion variables, using the same sets of covariates described previously.

## Results

Descriptive statistics for personality and intelligence measures are reported in Table 1.

**Table 1.**
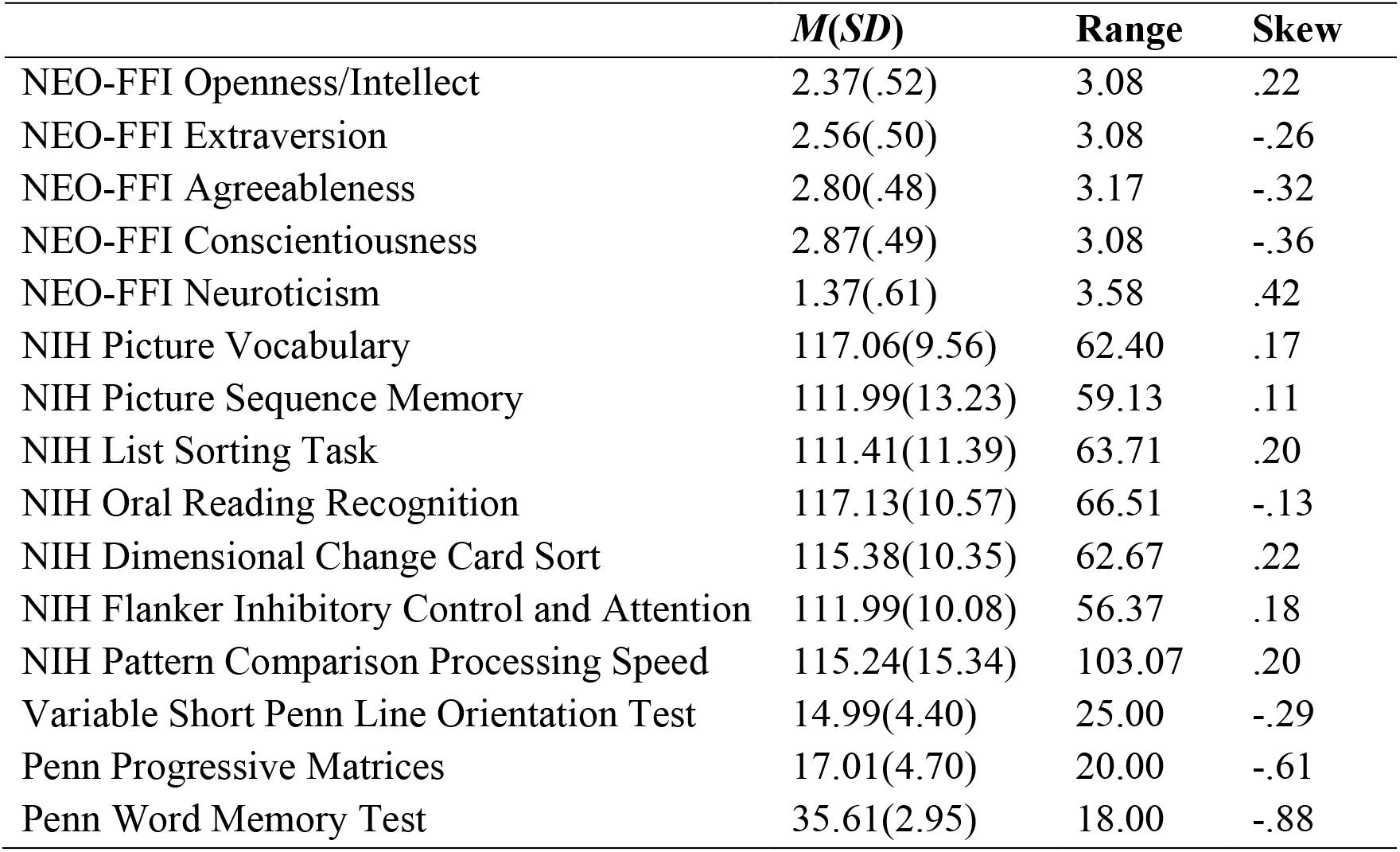
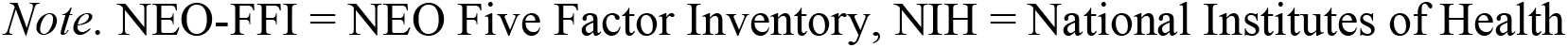
Descriptive statistics for personality and intelligence measures.

|  | *M(SD)* | **Range** | **Skew** |
| --- | --- | --- | --- |
| NEO-FFI Openness/Intellect | 2.37(.52) | 3.08 | .22 |
| NEO-FFI Extraversion | 2.56(.50) | 3.08 | -.26 |
| NEO-FFI Agreeableness | 2.80(.48) | 3.17 | -.32 |
| NEO-FFI Conscientiousness | 2.87(.49) | 3.08 | -.36 |
| NEO-FFI Neuroticism | 1.37(.61) | 3.58 | .42 |
| NIH Picture Vocabulary | 117.06(9.56) | 62.40 | .17 |
| NIH Picture Sequence Memory | 111.99(13.23) | 59.13 | .11 |
| NIH List Sorting Task | 111.41(11.39) | 63.71 | .20 |
| NIH Oral Reading Recognition | 117.13(10.57) | 66.51 | -.13 |
| NIH Dimensional Change Card Sort | 115.38(10.35) | 62.67 | .22 |
| NIH Flanker Inhibitory Control and Attention | 111.99(10.08) | 56.37 | .18 |
| NIH Pattern Comparison Processing Speed | 115.24(15.34) | 103.07 | .20 |
| Variable Short Penn Line Orientation Test | 14.99(4.40) | 25.00 | -.29 |
| Penn Progressive Matrices | 17.01(4.70) | 20.00 | -.61 |
| Penn Word Memory Test | 35.61(2.95) | 18.00 | -.88 |

*Note.* NEO-FFI = NEO Five Factor Inventory, NIH = National Institutes of Health

### Individual Parcel Flexibility

Before investigating associations of Openness/Intellect and intelligence with flexibility measures, we first evaluated the distribution of parcel flexibility measures across the cortex.

Parcels’ relative flexibility values remained largely consistent across partitions, with greater parcel flexibility observed in regions of the bilateral insula, medial frontal and parietal cortex, and reduced parcel flexibility in the lateral somatomotor and primary visual cortex, as well as regions of the inferior parietal lobule. Mean flexibility values were more variable across the cortex in the larger community partition, and more uniform in the partition with smaller communities. This pattern of parcel flexibility is similar to previous findings describing inter-subject functional connectivity variability patterns (Kong et al. 2019), and is largely consistent with previous descriptions of regions of cortex constituting a “flexible club” (Betzel, Satterthwaite et al. 2017; Yin et al. 2020). Mean values of overall flexibility for each parcel across the three community partitions are illustrated in Figure 2.

**Figure 2.**
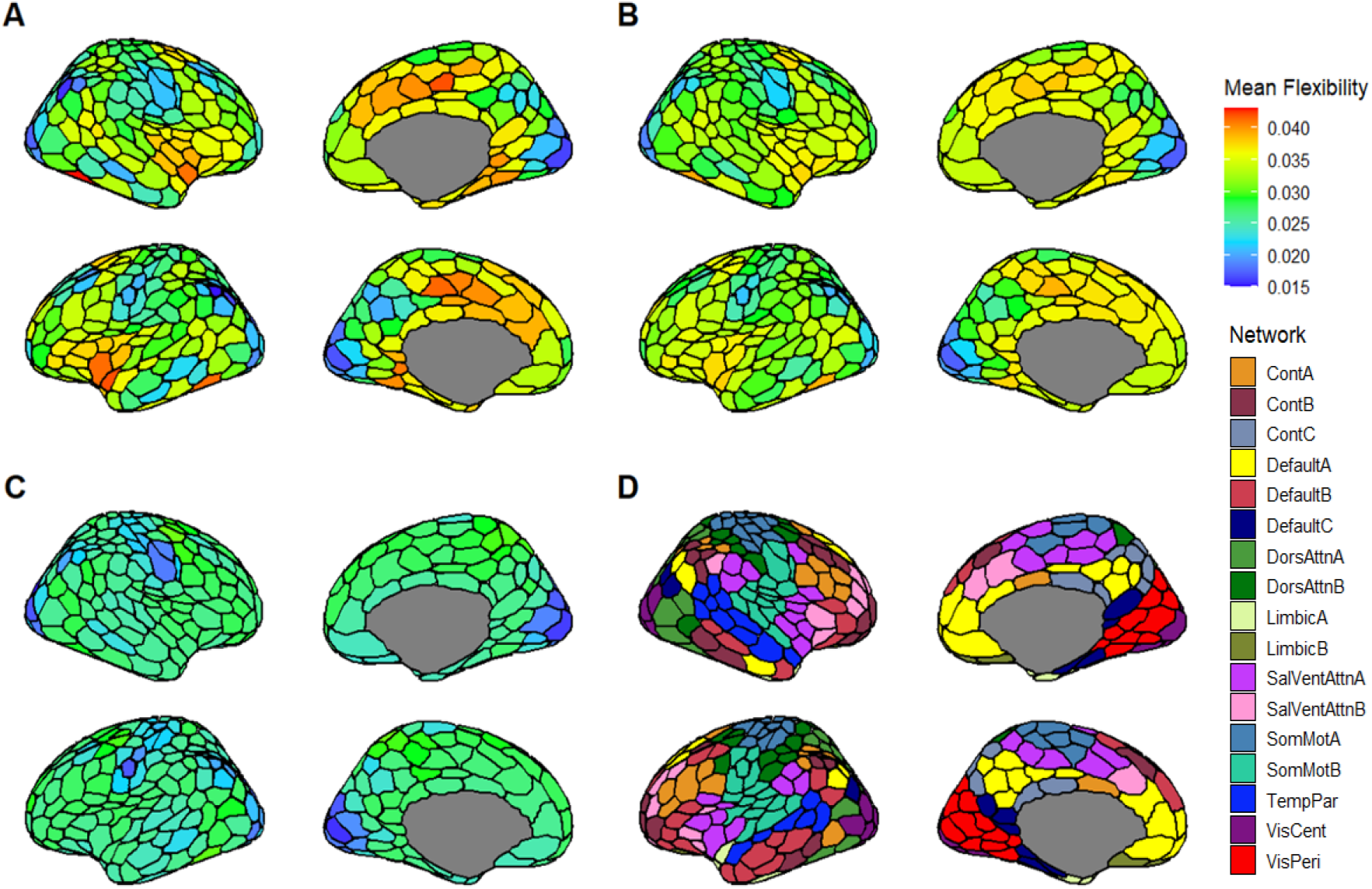
Mean overall flexibility across all participants derived from the communities optimized at γ = 1 (**A**), 1.1 (**B**), and 1.2 (**C**). Because each community partition reflects a different number of communities used to compute the flexibility of each parcel, mean flexibility values are not directly comparable across panels. Network assignments of parcels to 17 functional networks in the Schaefer et al. (2018) atlas are shown in panel **D**. Cont = frontoparietal control, DorsAttn = dorsal attention, SalVentAttn = salience/ventral attention, SomMot = somatomotor, TempPar = temporoparietal, VisCent = central visual, VisPeri = peripheral visual

Distributions of whole brain flexibility values for each community partition and flexibility type are illustrated in Figure 3. Cohesive and disjoint flexibility values differed across partitions as a result of the number and size of communities. At γ = 1, having only a few large communities resulted in a small absolute number of changes, most of which were cohesive, thus yielding disjoint flexibility values near zero. Conversely, at γ = 1.2, multiple small communities increased the likelihood of parcels changing affiliation independently, resulting in greater disjoint flexibility. Considering the relatively few number of functional communities and low disjoint flexibility values at the γ = 1 partition, we limited our hypothesis testing to the γ = 1.1 and 1.2 partitions to explore the unique predictive ability of cohesive and disjoint flexibility. (For completeness, we report analyses using γ = 1 in the online supplement.)

**Figure 3.**
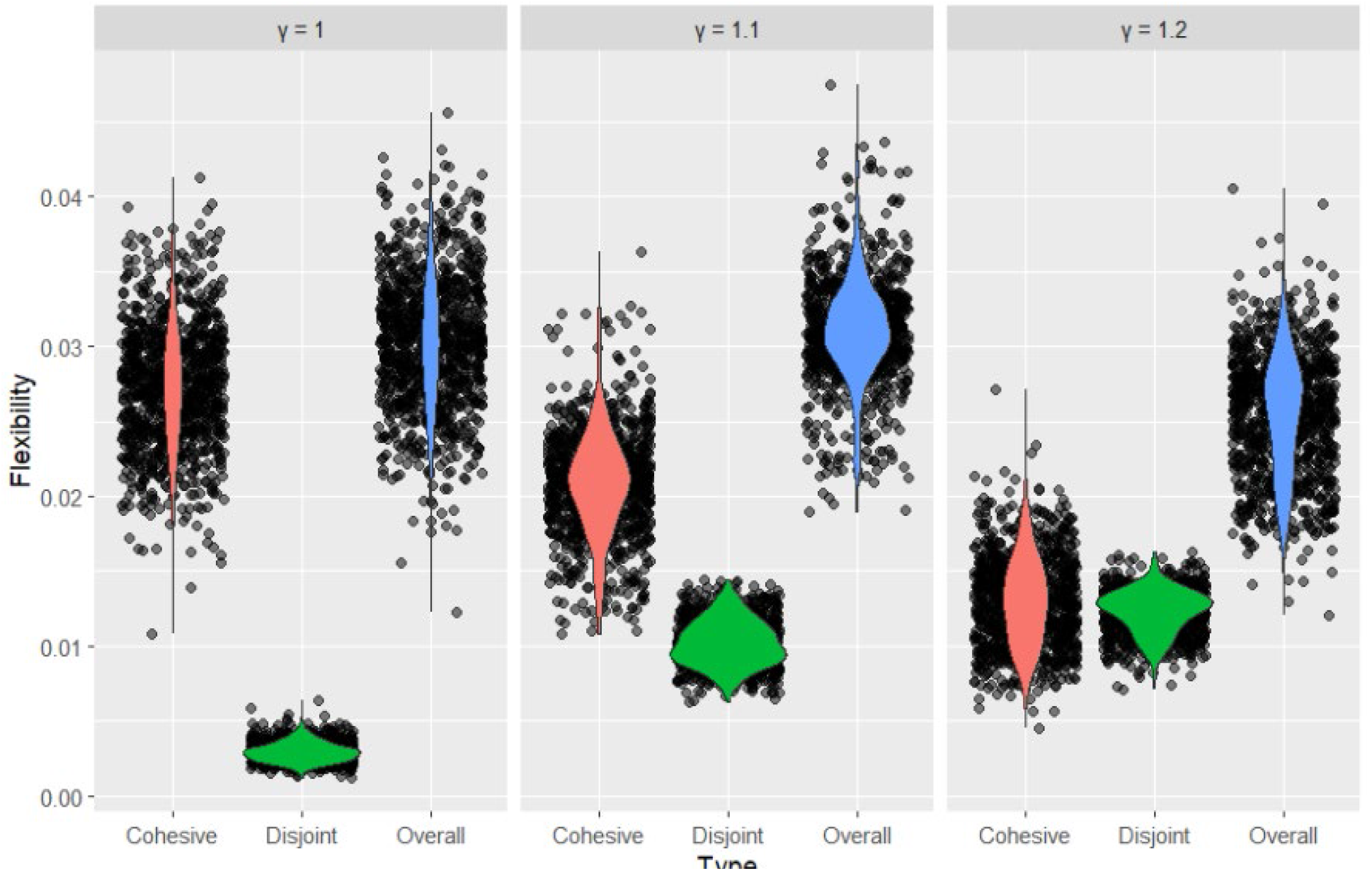
Distributions of whole brain flexibility values by flexibility type and community partition.

### Model Performance

Model parameters and performance metrics in the training and test sets for models using all parcels are reported in Table 2. Coefficients for all significant models are reported in the online supplement.

**Table 2.** Performance metrics of models using flexibility of all parcels as predictors.

| Criterion | Community | Type | Training Sample |  |  |  |  | Test Sample |  |  |
| --- | --- | --- | --- | --- | --- | --- | --- | --- | --- | --- |
| | | | $\alpha$ | $\lambda$ | RMSE | MAE | $R^2$ | RMSE | MAE | $R^2$ |
| O/I | $\gamma = 1.1$ | Overall | 0.8 | .0651691 | .99 | .78 | .047 | 1.04 | .83 | .000 |
|  |  | Cohesive | 0.2 | .0000621 | 2.58 | 2.10 | .040 | 2.57 | 2.04 | .001 |
|  |  | Disjoint | 0.6 | .1333822 | 1.00 | .78 | .041 | 1.01 | .81 | .001 |
| | $\gamma = 1.2$ | Overall | 0.7 | .0462009 | 1.00 | .80 | .060 | 1.04 | .84 | .010* |
|  |  | Cohesive | 0.4 | .0007842 | 2.58 | 2.05 | .055 | 2.35 | 1.87 | .002 |
|  |  | Disjoint | 0.8 | .0247086 | 1.12 | .89 | .026 | 1.17 | .93 | .000 |
| g | $\gamma = 1.1$ | Overall | 1 | .1016260 | 1.01 | .79 | .046 | 1.00 | .81 | .000 |
|  |  | Cohesive | 0.1 | .0000586 | 3.00 | 2.40 | .011 | 2.52 | 1.95 | .016** |
|  |  | Disjoint | 0.9 | .1050100 | 1.01 | .80 | .059 | .99 | .81 | .002 |
| | $\gamma = 1.2$ | Overall | 0.4 | .1272649 | 1.01 | .79 | .032 | 1.01 | .82 | .004 |
|  |  | Cohesive | 0.1 | .0001435 | 3.21 | 2.52 | .037 | 2.29 | 1.81 | .002 |
|  |  | Disjoint | 1 | .0003556 | 2.37 | 1.91 | .044 | 2.19 | 1.74 | .000 |

*Note.* \*permutation $p < .05$ , \*\*permutation $p < .01$ . g = general intelligence. O/I =
Openness/Intellect. $\alpha$ = mixing parameter, $\lambda$ = penalty parameter, RMSE = root-mean-square error. MAE = mean absolute error

#### Openness/Intellect

Openness/Intellect was not significantly associated with any flexibility measures in the γ = 1.1 community partition. In the communities identified at γ = 1.2, overall flexibility significantly predicted 1% of the variance in Openness/Intellect in the independent test set (permutation *p* = .03). Prediction of Openness/Intellect increased only slightly when removing intelligence as a covariate (*R*^2^ = .015, permutation *p* < .01), and remained unchanged when removing the rest of the Big Five (*R*^2^ = .01, permutation *p* = .03). The model included 136 parcels as predictors, distributed across all 17 networks, with the greatest frequency of parcels in the somatomotor, dorsal attention B, and default B networks. Openness/Intellect was most strongly associated with increased flexibility in the bilateral extrastriate cortex, postcentral gyrus, and dorsal prefrontal cortex, and with reduced flexibility in regions of the somatomotor cortex, ventrolateral prefrontal cortex, and frontal operculum. However, results from a Kruskal-Wallis rank sum test indicated no significant differences in median parcel coefficient magnitude across networks (*H*(16) = 9.8747, *p* = .87). Model coefficients are illustrated in Figure 4.

**Figure 4.**
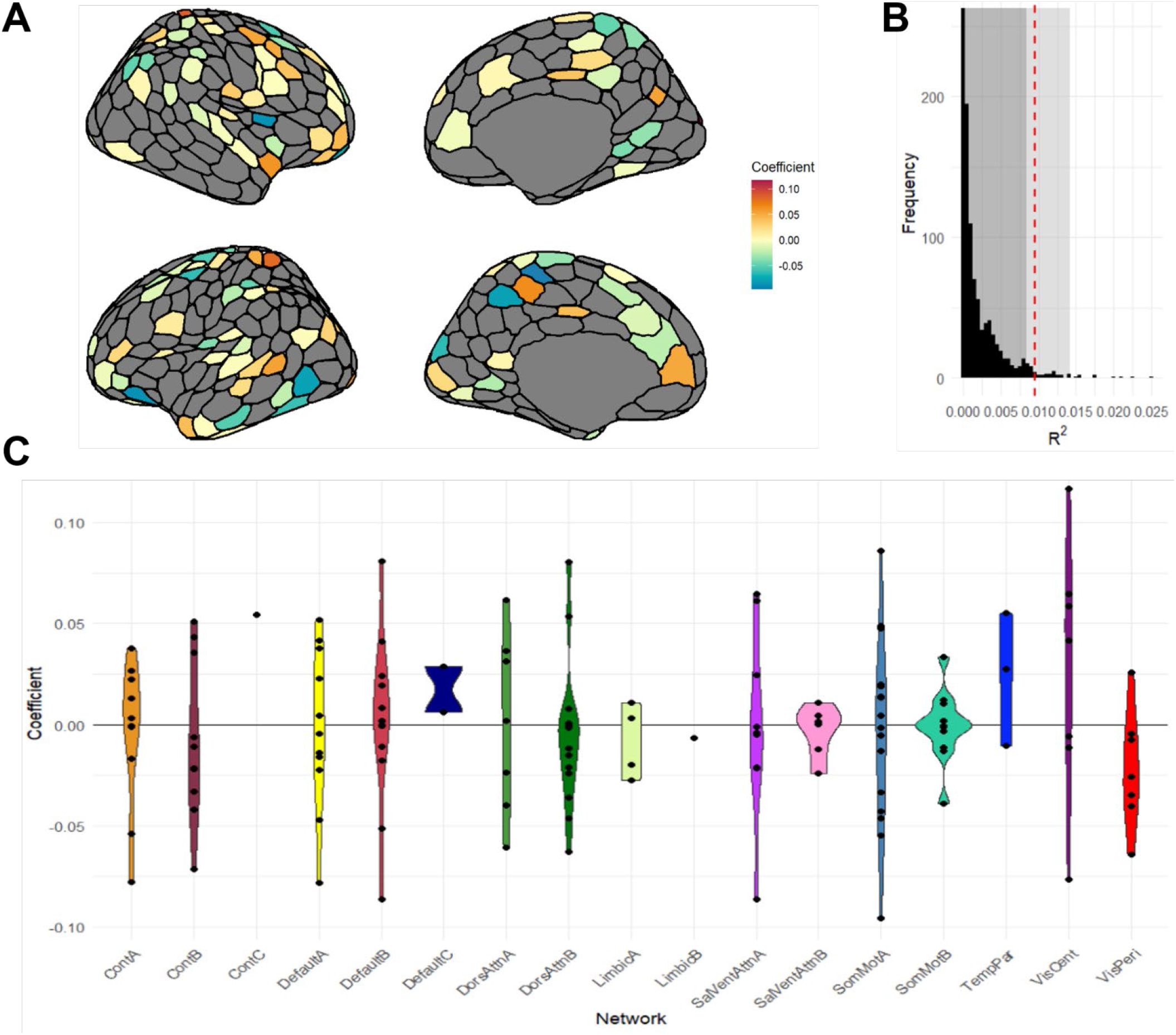
Model predicting Openness/Intellect from overall parcel flexibility. **A**: Model coefficients selected in the community partition optimized at γ = 1.2. Parcels not retained in the model are shown in gray. **B:** Null distribution of permuted *R*^2^ values. The red dashed line indicates the observed *R*^2^ value. The dark shaded region signifies values at *p* ≥ .05. The light shaded region signifies values at *p* ≥ .01. **C:** Violin plots of model coefficients sorted by 17 functional networks from the Schaefer et al. (2018) atlas

#### Intelligence

Intelligence was not significantly associated with any flexibility measures in the γ = 1.2 community partition. In the communities identified at γ = 1.1, cohesive flexibility significantly predicted 1.6% of the variance in intelligence in the independent test set (permutation *p* < .01). Cohesive flexibility also significantly predicted intelligence when including the Big Five as covariates (*R*^2^ = .009, permutation *p* = .03). As with Openness/Intellect, this model included parcels from all networks, but in contrast included all 400 parcels as predictors. Intelligence was most strongly associated with increased cohesive flexibility of parcels in the bilateral extrastriate and somatomotor cortex, superior parietal lobule, frontal pole, and dorsolateral prefrontal cortex (dlPFC), as well as decreased flexibility in regions of the primary visual cortex, postcentral gyrus, and precuneus, but results from a Kruskal-Wallis rank sum test again indicated no significant differences in median parcel coefficient magnitude across networks (*H*(16) = 7.3246, *p* = .97). Parcels included in this model are illustrated in Figure 5.

**Figure 5.**
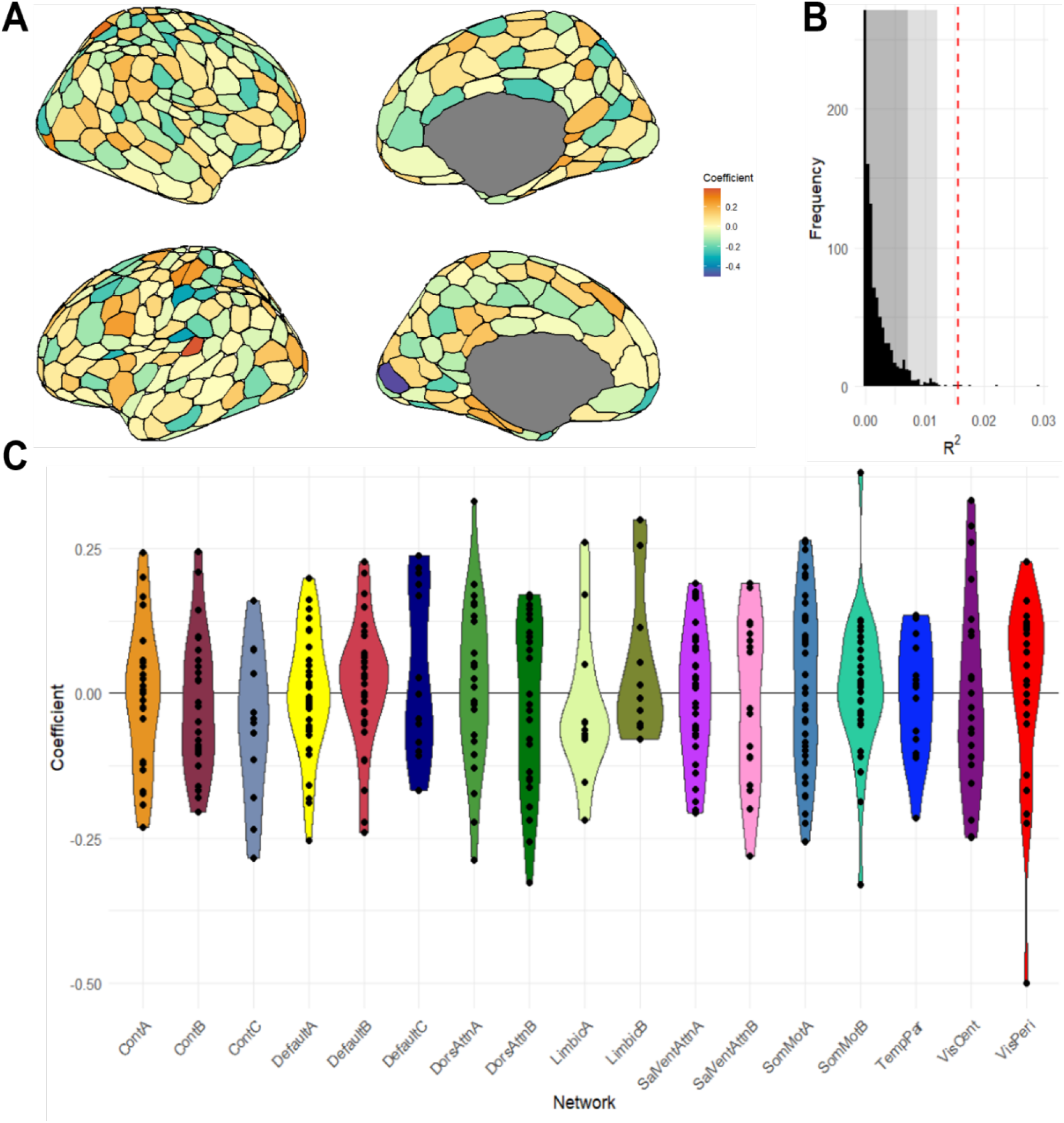
Model predicting intelligence from parcel cohesive flexibility. **A**: Model coefficients selected in the community partition optimized at γ = 1.1. **B:** Null distribution of permuted *R*^2^ values. The red dashed line indicates the observed *R*^2^ value. The dark shaded region signifies values at *p* ≥ .05. The light shaded region signifies values at *p* ≥ .01. **C:** Violin plots of model coefficients sorted by 17 functional networks from the Schaefer et al. (2018) atlas.

#### Discriminant Validity

Additional tests of discriminant validity revealed that all flexibility types were not significantly associated with any of the remaining Big Five across all community partitions. Performance metrics of models predicting the remaining Big Five are included in Table 3.

**Table 3.**
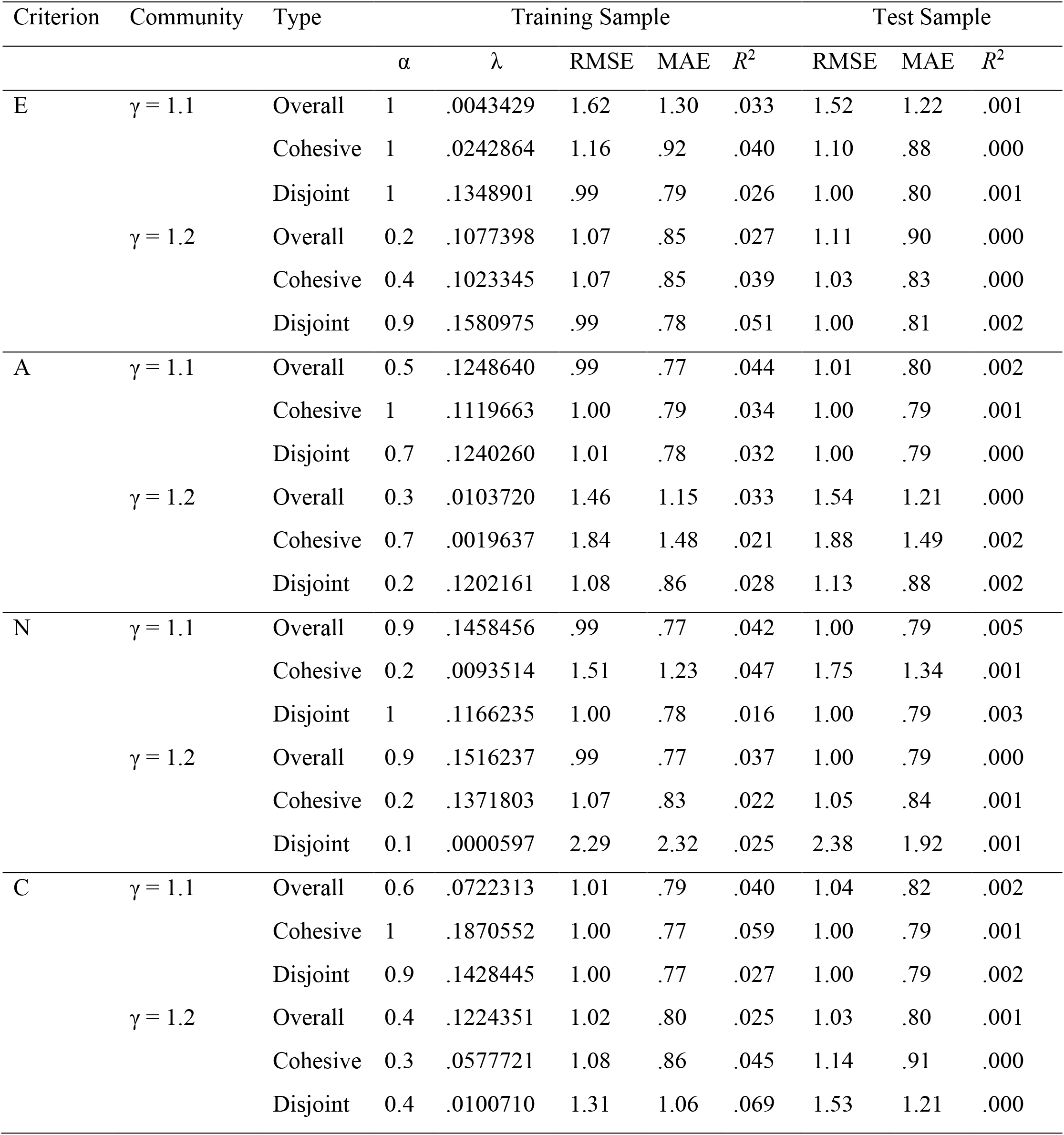
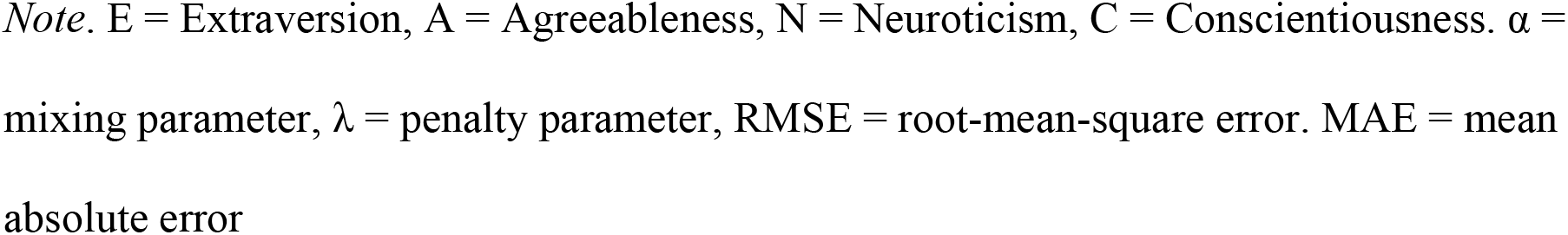
Performance metrics of models predicting the remaining Big Five from parcel flexibility.

| Criterion | Community | Type | Training Sample |  |  |  |  | Test Sample |  |  |
| --- | --- | --- | --- | --- | --- | --- | --- | --- | --- | --- |
| | | | $\alpha$ | $\lambda$ | RMSE | MAE | $R^2$ | RMSE | MAE | $R^2$ |
| E | $\gamma = 1.1$ | Overall | 1 | .0043429 | 1.62 | 1.30 | .033 | 1.52 | 1.22 | .001 |
|  |  | Cohesive | 1 | .0242864 | 1.16 | .92 | .040 | 1.10 | .88 | .000 |
|  |  | Disjoint | 1 | .1348901 | .99 | .79 | .026 | 1.00 | .80 | .001 |
| | $\gamma = 1.2$ | Overall | 0.2 | .1077398 | 1.07 | .85 | .027 | 1.11 | .90 | .000 |
|  |  | Cohesive | 0.4 | .1023345 | 1.07 | .85 | .039 | 1.03 | .83 | .000 |
|  |  | Disjoint | 0.9 | .1580975 | .99 | .78 | .051 | 1.00 | .81 | .002 |
| A | $\gamma = 1.1$ | Overall | 0.5 | .1248640 | .99 | .77 | .044 | 1.01 | .80 | .002 |
|  |  | Cohesive | 1 | .1119663 | 1.00 | .79 | .034 | 1.00 | .79 | .001 |
|  |  | Disjoint | 0.7 | .1240260 | 1.01 | .78 | .032 | 1.00 | .79 | .000 |
| | $\gamma = 1.2$ | Overall | 0.3 | .0103720 | 1.46 | 1.15 | .033 | 1.54 | 1.21 | .000 |
|  |  | Cohesive | 0.7 | .0019637 | 1.84 | 1.48 | .021 | 1.88 | 1.49 | .002 |
|  |  | Disjoint | 0.2 | .1202161 | 1.08 | .86 | .028 | 1.13 | .88 | .002 |
| N | $\gamma = 1.1$ | Overall | 0.9 | .1458456 | .99 | .77 | .042 | 1.00 | .79 | .005 |
|  |  | Cohesive | 0.2 | .0093514 | 1.51 | 1.23 | .047 | 1.75 | 1.34 | .001 |
|  |  | Disjoint | 1 | .1166235 | 1.00 | .78 | .016 | 1.00 | .79 | .003 |
| | $\gamma = 1.2$ | Overall | 0.9 | .1516237 | .99 | .77 | .037 | 1.00 | .79 | .000 |
|  |  | Cohesive | 0.2 | .1371803 | 1.07 | .83 | .022 | 1.05 | .84 | .001 |
|  |  | Disjoint | 0.1 | .0000597 | 2.29 | 2.32 | .025 | 2.38 | 1.92 | .001 |
| C | $\gamma = 1.1$ | Overall | 0.6 | .0722313 | 1.01 | .79 | .040 | 1.04 | .82 | .002 |
|  |  | Cohesive | 1 | .1870552 | 1.00 | .77 | .059 | 1.00 | .79 | .001 |
|  |  | Disjoint | 0.9 | .1428445 | 1.00 | .77 | .027 | 1.00 | .79 | .002 |
| | $\gamma = 1.2$ | Overall | 0.4 | .1224351 | 1.02 | .80 | .025 | 1.03 | .80 | .001 |
|  |  | Cohesive | 0.3 | .0577721 | 1.08 | .86 | .045 | 1.14 | .91 | .000 |
|  |  | Disjoint | 0.4 | .0100710 | 1.31 | 1.06 | .069 | 1.53 | 1.21 | .000 |

*Note.* E = Extraversion, A = Agreeableness, N = Neuroticism, C = Conscientiousness. $\alpha$ = mixing parameter, $\lambda$ = penalty parameter, RMSE = root-mean-square error. MAE = mean absolute error

## Discussion

Openness/Intellect, measured by questionnaire, and intelligence, measured by cognitive performance tests, were significantly associated with brain network flexibility. The prediction accuracy metrics reported in this study are small, but they accord with previous research using machine learning approaches with common functional connectivity measures to predict traits like Plasticity and Openness/Intellect (Dubois et al. 2018b), psychotic-like experiences (Ma et al. 2022), and executive function (Heckner et al. 2023). These results suggest that measures of flexibility in a dynamic functional connectivity framework may provide a similar degree of information about personality as static resting-state functional connectivity, using cross-validation procedures, despite recent evidence suggesting that dynamic and static functional connectivity describe different properties of neural activity (Zhang et al. 2023; Zhu et al. 2021).

Although the amount of variance explained by functional community flexibility is small, this result is nonetheless an important step in a larger effort to understand the neural correlates of broad personality domains as dynamical systems. Previous research has described associations of a variety of influential behavioral phenotypes like need for cognition, working memory, and creativity with flexibility (Braun et al. 2015; He et al. 2019), and these different traits and abilities are all situated within the larger Openness/Intellect domain (DeYoung et al. 2012; Saucier 1992). This research helps expand on these findings by demonstrating that brain network flexibility similarly predicts the broader Openness/Intellect domain beyond specific facets describing features of higher-order cognition.

Features of these results cohere with results of prior research predicting Openness/Intellect and intelligence from functional connectivity. For example, much research using static functional connectivity suggests that effect sizes for intelligence are larger than those for Openness/Intellect and other questionnaire variables (Kong et al. 2019; Li et al. 2019; Kong et al. 2021; Chen et al. 2022). One simple explanation for this pattern may be the fact that intelligence was assessed using a variety of different task measures, while Openness/Intellect was measured through a single, short self-report questionnaire. However, better measurement may not be the only reason for better prediction; there may be a more substantive reason as well, reflected in the fact that Openness/Intellect also seems to be more readily predicted from functional connectivity in the cortex than the other Big Five traits (Dubois et al. 2018b; Kong et al. 2021). Considering the conceptual similarity of intelligence and Openness/Intellect in describing variation in complex cognition, one might expect complex cognition to be reflected more widely across the cortex compared to the more specific affective and motivational functions underlying other traits.

We observed that the cohesive flexibility of all 400 parcels contributed to the best prediction of intelligence in the larger (γ = 1.1) community partition. This observation is consistent with the hypothesis by Safron et al. (2022) that cohesive flexibility indexes adaptive functioning. The fact that cohesive flexibility specifically predicts intelligence aligns with previous research suggesting that intelligence reflects a form of global network integrity (Langer et al. 2012; Barbey 2018; Wang et al. 2021), with intelligence predicting a variety of positive life outcomes (Gottfredson 1997; Brown et al. 2021). Although no significant differences were observed across networks, many of the parcels with the strongest effect sizes were located in cortical hubs nominated by the Parieto-Frontal Integration Theory (P-FIT; Jung & Haier 2007; Gur et al. 2021), including the dlPFC and superior parietal lobule.

These findings reveal that intelligence is associated with the tendency for networks to reorganize themselves frequently, throughout the cortex. Previous research has demonstrated the utility of using static functional connectivity profiles across the entire cortex to predict intelligence (Feilong et al. 2021), and our results indicate that a similar pattern exists using dynamic functional connectivity. The association of intelligence with cohesive flexibility across the cortex provides further evidence to support the notion that intelligence can be understood through integrative functional dynamics across the whole brain, but may be orchestrated by certain regions most implicated in higher-order cognition (Alavash et al. 2015; Avery et al. 2020; Cole et al. 2012; Hilger et al. 2017).

Despite its empirical and conceptual relation to intelligence, Openness/Intellect exhibited different patterns of associations with flexibility, and these remained significant even when controlling for intelligence. Rather than whole brain cohesive flexibility, Openness/Intellect was best predicted by the overall flexibility of a more limited set of parcels, in the smaller community partition (γ = 1.2), with the strongest effects observed in both somatosensory regions and areas implicated in abstract and self-referential thought like the dorsal prefrontal cortex and precuneus. Of note, Openness/Intellect was reliably predicted by overall flexibility specifically, rather than cohesive flexibility. Overall flexibility reflects both cohesive and disjoint flexibility. Safron et al. (2022) hypothesized that disjoint flexibility may be a marker of a general tendency toward neural entropy, and additional research has shown that this may be the case especially in regions involved in higher-order cognition (Telesford et al. 2017). Given that Openness/Intellect domain subsumes a variety of facets capturing elements of both adaptive and disorganized cognition (DeYoung et al. 2012), both cohesive and disjoint brain dynamics could be contributing unique variance to the prediction of this broad personality domain, especially when controlling for the effect of intelligence.

Previous research has implicated dopaminergic function in Openness/Intellect (DeYoung 2013; Passamonti et al. 2015), and D2 receptor binding specifically has been linked to cognitive flexibility (Durstewitz and Seamans 2008; Wacker et al. 2012). To connect our results with the hypothesized link between Openness/Intellect and dopamine, we conducted additional exploratory analyses using D2 receptor density positron emission tomography (PET) maps from Smith et al. (2019) and Sandiego et al. (2015), which were averaged and parcellated using the Schaefer atlas by Hansen et al. (2022). These receptor maps are available at https://github.com/netneurolab/hansen_receptors. We found that D2 receptor density was significantly positively associated with overall flexibility across all parcels in the γ = 1.2 partition (*r* = .11, *p* < .05), the same partition associated with Openness/Intellect, but not with flexibility in the γ = 1.1 (*r* = .06, *p* = .20) or γ = 1 partitions (*r* = .02, *p* = .68).

This association might suggest a potential neurobiological mechanism underlying the kind of community reconfiguration that contributes to cognitive flexibility. D2 dopamine neurons projecting to prefrontal cortical regions from midbrain structures are described as salience coding neurons, and support cognitive processes relating to exploration of information (Bromberg-Martin et al., 2010). This engagement with complex information can involve multiple different cognitive operations, including the redirection of attentional resources, the manipulation of information in working memory, self-referential thought, and the management of different or conflicting goal states. Further, tasks involving complex cognition have been shown to recruit multiple cortical networks with hubs in prefrontal regions, often at different times during the task (Beaty et al. 2015; Patil et al. 2021). It may be that cortical regions with greater D2 receptor density participate in more frequent but relatively localized community changes as a means of flexibly managing the cognitive demand of identifying and exploring abstract or sensory information.

### Flexibility and Individualized Parcellation

One particular strength of our method is the use of individualized parcellation. Previous research has described a variety of properties differing between group-level atlases and their individualized counterparts, with the latter yielding increased effect sizes in brain-behavior associations (Kong et al. 2021; Sassenberg et al. 2023) and more conservative patterns of functional connectivity in case-control designs (Levi et al. 2023). To the best of our knowledge, no other research investigating brain-behavior associations with flexibility has made use of individualized parcellation. Other research using similar procedures to identify functional communities, but without individualized parcellation, has reported similar but slightly larger flexibility values (Yin et al. 2020). It may be that more accurate localization of parcel boundaries, using individualized parcellation, produces a more conservative but more accurate estimation of community dynamics over time. Parcel boundaries in group-level atlases do not respect the individual variation in functional topography on the cortical surface, and so canonical parcels without individualization may instead capture distinct functional regions within the same boundary, leading to estimates of greater flexibility than what actually occurs. For instance, a poorly estimated parcel boundary could erroneously group two functional regions belonging to networks with negatively correlated activation patterns like the default and frontoparietal control networks. That parcel might then exhibit greater flexibility as it appears to change affiliation with other communities containing default or frontoparietal control network parcels, simply depending on the relative activation patterns of each network in a given window.

### Limitations

The present study incorporated a number of cutting-edge analytic strategies beyond individualized parcellation, including calculation of functional connectivity through magnitude squared coherence and the use of the 400-parcel Schaefer atlas. Evidence suggests that magnitude squared coherence performs at least as well as product-moment correlations as a measure of functional connectivity (Betzel, Satterthwaite et al. 2017) and that the Schaefer atlas exhibits greater within-parcel homogeneity compared to its competitors (Schaefer et al. 2018). However, there are also important limitations to consider. One limitation is that these analyses included flexibility metrics only for cortical parcels. Examination of subcortical structures involved in dopamine modulation might provide additional insight in relating Openness/Intellect and intelligence to brain network flexibility.

Additionally, the particular parameters of the multi-layer modularity algorithm in the present analyses were chosen to explore results across different spatial resolutions of the cortical communities, but we did not explore different temporal resolutions when calculating network flexibility. Fixing the ⍵ parameter to 1 for all analyses was done to manage the computational intensity and number of analyses, but that means we do not know how results might be affected by employing different temporal windows in our analysis of flexibility.

Further, this study is limited by the use of a sliding-window approach to dynamic functional connectivity. Future research can explore whether these associations persist using more sophisticated measures of functional connectivity dynamics like dwell time. Our measure of flexibility also may be limited by the features of the general Louvain community optimization algorithm, which categorizes parcels into discrete communities. Explorations into more nuanced measures of community partitioning like stochastic block modeling might also improve prediction accuracy (Lee and Wilkinson 2019).

The relatively low prediction accuracies derived from the current findings also echo recent concerns of whether the explained variance is useful for identifying precise neural mechanisms underlying these traits, as well as whether these performance metrics would improve substantially through the use of task-based designs (Finn 2021; DeYoung et al. 2022). Further research would benefit from examining model performance under conditions where participants are engaged in tasks that elicit cognitive states corresponding to particular traits of interest, as it may be the case that dynamic reconfigurations of cortical topology are more pronounced (or importantly different in other ways) from dynamics observed at rest (Greene et al. 2018; Rzucidlo et al. 2013; Salehi et al. 2020).

Lastly, future research exploring associations of cortical flexibility with measures of personality would benefit from improved measurement of behavioral constructs. Although a measure of intelligence in the present research was constructed using a variety of different tasks, Openness/Intellect was limited to a single, relatively short questionnaire measure. Future research could use multiple self- and peer-report measures of personality and use latent variable analyses to model personality constructs.

## Conclusion

This research presents the first empirical test of a hypothesis relating neural flexibility to broad personality traits describing cognitive and behavioral flexibility, inspired by the work of Safron et al. (2022). We describe reliable associations between intelligence and a pattern of cohesive flexibility across the entire cortex, as well as between Openness/Intellect and a more delimited pattern of general flexibility. These findings help to expand on previous research describing associations of neural flexibility with a variety of behavioral and cognitive traits relating to higher-order cognition within the broader framework of the personality hierarchy. This research also provides further support for previous research reporting associations between functional connectivity and these traits in the Human Connectome Project, as well as helping to frame the neural correlates of these traits as dynamical systems.

## Acknowledgements

We would like to thank Victoria Klimaj for her feedback on this work. We also thank the collectors and curators of the data from the Human Connectome Project.

