## Supplementary material for "Stable Individual Differences from Dynamic Patterns of Function: Brain Network Flexibility Predicts Openness/Intellect and Intelligence": Online Supplement

This supplementary material contains tables of model coefficients, additional model performance metrics in larger functional communities, and results of models using network flexibility to predict metatrait Plasticity as a manifest variable.

### Table of Contents

Coefficients of models predicting intelligence and Openness/Intellect 2

Performance metrics of models in the $\gamma = 1$ community partition 13

Predicting metatrait Plasticity from network flexibility 14

Supplementary references 15

**Table A1**

*Coefficients of models predicting intelligence and Openness/Intellect from parcel flexibility*

| Parcel Index | Parcel Name | *g* cohesive, γ = 1.1 | O/I overall, γ = 1.2 |
|---|---|---|---|
| 1 | 17Networks_LH_VisCent_ExStr_1 | -0.155293109 | -0.011185613 |
| 2 | 17Networks_LH_VisCent_ExStr_2 | -0.219417318 | |
| 3 | 17Networks_LH_VisCent_ExStr_3 | -0.122474174 | |
| 4 | 17Networks_LH_VisCent_ExStr_4 | 0.026471456 | |
| 5 | 17Networks_LH_VisCent_ExStr_5 | 0.099852645 | 0.058841811 |
| 6 | 17Networks_LH_VisCent_ExStr_6 | -0.06850162 | |
| 7 | 17Networks_LH_VisCent_Striate_1 | -0.108161795 | |
| 8 | 17Networks_LH_VisCent_ExStr_7 | 0.029720221 | |
| 9 | 17Networks_LH_VisCent_ExStr_8 | 0.260804201 | |
| 10 | 17Networks_LH_VisCent_ExStr_9 | 0.197596191 | |
| 11 | 17Networks_LH_VisCent_ExStr_10 | 0.128293487 | 0.064724079 |
| 12 | 17Networks_LH_VisCent_ExStr_11 | 0.00160869 | |
| 13 | 17Networks_LH_VisPeri_ExStrInf_1 | 0.12239046 | |
| 14 | 17Networks_LH_VisPeri_ExStrInf_2 | 0.111246392 | |
| 15 | 17Networks_LH_VisPeri_ExStrInf_3 | -0.037092588 | -0.00733969 |
| 16 | 17Networks_LH_VisPeri_ExStrInf_4 | -0.0000456 | |
| 17 | 17Networks_LH_VisPeri_ExStrInf_5 | -0.208359695 | -0.02580127 |
| 18 | 17Networks_LH_VisPeri_StriCal_1 | -0.500687879 | 0.025623399 |
| 19 | 17Networks_LH_VisPeri_StriCal_2 | 0.115168823 | |
| 20 | 17Networks_LH_VisPeri_ExStrSup_1 | 0.084984731 | |
| 21 | 17Networks_LH_VisPeri_ExStrSup_2 | 0.120600476 | -0.064290309 |
| 22 | 17Networks_LH_VisPeri_ExStrSup_3 | -0.140365486 | |
| 23 | 17Networks_LH_VisPeri_ExStrSup_4 | 0.160654653 | |
| 24 | 17Networks_LH_VisPeri_ExStrSup_5 | 0.048634443 | |
| 25 | 17Networks_LH_SomMotA_1 | -0.091098518 | |
| 26 | 17Networks_LH_SomMotA_2 | 0.087515476 | |
| 27 | 17Networks_LH_SomMotA_3 | 0.265371529 | |
| 28 | 17Networks_LH_SomMotA_4 | 0.249295024 | |
| 29 | 17Networks_LH_SomMotA_5 | 0.032792507 | |
| 30 | 17Networks_LH_SomMotA_6 | -0.0211299 | -0.095797256 |
| 31 | 17Networks_LH_SomMotA_7 | 0.088709537 | |
| 32 | 17Networks_LH_SomMotA_8 | 0.134182425 | |
| 33 | 17Networks_LH_SomMotA_9 | -0.04608867 | 0.013301129 |
| 34 | 17Networks_LH_SomMotA_10 | 0.069271948 | |
| 35 | 17Networks_LH_SomMotA_11 | -0.175416083 | 0.048970197 |
| 36 | 17Networks_LH_SomMotA_12 | 0.218968157 | -0.05461564 |
| 37 | 17Networks_LH_SomMotA_13 | -0.047982939 | -0.04288802 |
| 38 | 17Networks_LH_SomMotA_14 | 0.201707832 | -0.00515708 |
| 39 | 17Networks_LH_SomMotA_15 | -0.069590704 | -0.033530349 |
| 40 | 17Networks_LH_SomMotA_16 | 0.26203595 | 0.004502853 |
| 41 | 17Networks_LH_SomMotA_17 | 0.010011555 | |

| | | | |
|---|---|---|---|
| 42 | 17Networks_LH_SomMotA_18 | -0.155782671 | |
| 43 | 17Networks_LH_SomMotA_19 | 0.156301207 | -0.054957611 |
| 44 | 17Networks_LH_SomMotB_Aud_1 | -0.013058687 | |
| 45 | 17Networks_LH_SomMotB_Aud_2 | 0.018445774 | -0.001042239 |
| 46 | 17Networks_LH_SomMotB_Ins_1 | 0.126081231 | -0.001153091 |
| 47 | 17Networks_LH_SomMotB_S2_1 | 0.021239578 | |
| 48 | 17Networks_LH_SomMotB_S2_2 | -0.033269697 | |
| 49 | 17Networks_LH_SomMotB_Aud_3 | -0.036026114 | |
| 50 | 17Networks_LH_SomMotB_Aud_4 | 0.382400135 | -0.012882738 |
| 51 | 17Networks_LH_SomMotB_S2_3 | 0.036641245 | -0.03896458 |
| 52 | 17Networks_LH_SomMotB_S2_4 | 0.011671331 | |
| 53 | 17Networks_LH_SomMotB_S2_5 | -0.330930386 | 0.002087179 |
| 54 | 17Networks_LH_SomMotB_S2_6 | 0.089329714 | |
| 55 | 17Networks_LH_SomMotB_Cent_1 | 0.015184432 | |
| 56 | 17Networks_LH_SomMotB_Cent_2 | -0.031231944 | |
| 57 | 17Networks_LH_SomMotB_Cent_3 | 0.116709626 | |
| 58 | 17Networks_LH_SomMotB_Cent_4 | 0.014703624 | |
| 59 | 17Networks_LH_SomMotB_Cent_5 | 0.002103135 | |
| 60 | 17Networks_LH_DorsAttnA_TempOcc_1 | -0.071373337 | 0.061575427 |
| 61 | 17Networks_LH_DorsAttnA_TempOcc_2 | -0.105952222 | |
| 62 | 17Networks_LH_DorsAttnA_TempOcc_3 | -0.172313559 | -0.060545889 |
| 63 | 17Networks_LH_DorsAttnA_TempOcc_4 | 0.133281199 | |
| 64 | 17Networks_LH_DorsAttnA_ParOcc_1 | 0.156642316 | |
| 65 | 17Networks_LH_DorsAttnA_ParOcc_2 | 0.046846647 | 0.03662701 |
| 66 | 17Networks_LH_DorsAttnA_SPL_1 | -0.082208802 | |
| 67 | 17Networks_LH_DorsAttnA_SPL_2 | 0.18831794 | |
| 68 | 17Networks_LH_DorsAttnA_SPL_3 | 0.070612272 | |
| 69 | 17Networks_LH_DorsAttnA_SPL_4 | 0.169601905 | |
| 70 | 17Networks_LH_DorsAttnA_SPL_5 | -0.082476396 | |
| 71 | 17Networks_LH_DorsAttnA_SPL_6 | 0.069500249 | |
| 72 | 17Networks_LH_DorsAttnA_SPL_7 | -0.287785257 | |
| 73 | 17Networks_LH_DorsAttnB_PostC_1 | -0.018536582 | |
| 74 | 17Networks_LH_DorsAttnB_PostC_2 | -0.326799032 | -0.024149038 |
| 75 | 17Networks_LH_DorsAttnB_PostC_3 | -0.255626226 | -0.001493467 |
| 76 | 17Networks_LH_DorsAttnB_PostC_4 | -0.079096683 | -0.046067399 |
| 77 | 17Networks_LH_DorsAttnB_PostC_5 | -0.161547534 | |
| 78 | 17Networks_LH_DorsAttnB_PostC_6 | -0.001676372 | 0.080444634 |
| 79 | 17Networks_LH_DorsAttnB_PostC_7 | 0.089477347 | 0.007929896 |
| 80 | 17Networks_LH_DorsAttnB_PostC_8 | -0.08840555 | |
| 81 | 17Networks_LH_DorsAttnB_PostC_9 | -0.029538171 | 0.000843178 |
| 82 | 17Networks_LH_DorsAttnB_FEF_1 | 0.060775928 | -0.062696361 |
| 83 | 17Networks_LH_DorsAttnB_FEF_2 | -0.134758144 | -0.011589896 |
| 84 | 17Networks_LH_DorsAttnB_FEF_3 | 0.075287678 | 0.000426864 |
| 85 | 17Networks_LH_DorsAttnB_PrCv_1 | 0.146313749 | |

| | | | |
|---|---|---|---|
| 86 | 17Networks_LH_SalVentAttnA_ParOper_1 | -0.186439461 | |
| 87 | 17Networks_LH_SalVentAttnA_ParOper_2 | -0.062711044 | |
| 88 | 17Networks_LH_SalVentAttnA_ParOper_3 | 0.074412527 | |
| 89 | 17Networks_LH_SalVentAttnA_Ins_1 | 0.189950476 | |
| 90 | 17Networks_LH_SalVentAttnA_Ins_2 | -0.123328433 | |
| 91 | 17Networks_LH_SalVentAttnA_Ins_3 | 0.09111905 | |
| 92 | 17Networks_LH_SalVentAttnA_Ins_4 | 0.171222004 | |
| 93 | 17Networks_LH_SalVentAttnA_FrOper_1 | 0.015436748 | |
| 94 | 17Networks_LH_SalVentAttnA_FrOper_2 | -0.061818394 | |
| 95 | 17Networks_LH_SalVentAttnA_ParMed_1 | -0.137971564 | |
| 96 | 17Networks_LH_SalVentAttnA_ParMed_2 | -0.056054134 | 0.064782843 |
| 97 | 17Networks_LH_SalVentAttnA_ParMed_3 | 0.01660256 | |
| 98 | 17Networks_LH_SalVentAttnA_FrMed_1 | -0.123180703 | |
| 99 | 17Networks_LH_SalVentAttnA_FrMed_2 | 0.122290263 | -0.021442034 |
| 100 | 17Networks_LH_SalVentAttnA_FrMed_3 | -0.06912153 | |
| 101 | 17Networks_LH_SalVentAttnB_PFCl_1 | -0.159350386 | |
| 102 | 17Networks_LH_SalVentAttnB_PFCl_2 | 0.071244482 | |
| 103 | 17Networks_LH_SalVentAttnB_PFCl_3 | 0.103651632 | |
| 104 | 17Networks_LH_SalVentAttnB_Ins_1 | -0.034146675 | |
| 105 | 17Networks_LH_SalVentAttnB_Ins_2 | -0.167206363 | |
| 106 | 17Networks_LH_SalVentAttnB_Ins_3 | -0.280475376 | 0.010767169 |
| 107 | 17Networks_LH_SalVentAttnB_OFC_1 | 0.119577059 | |
| 108 | 17Networks_LH_SalVentAttnB_PFCmp_1 | -0.110896905 | -0.012097999 |
| 109 | 17Networks_LH_LimbicB_OFC_1 | 0.01420766 | |
| 110 | 17Networks_LH_LimbicB_OFC_2 | -0.079714712 | -0.006706223 |
| 111 | 17Networks_LH_LimbicB_OFC_3 | -0.058368476 | |
| 112 | 17Networks_LH_LimbicB_OFC_4 | 0.053595146 | |
| 113 | 17Networks_LH_LimbicB_OFC_5 | 0.300330639 | |
| 114 | 17Networks_LH_LimbicA_TempPole_1 | -0.15246149 | 0.011064964 |
| 115 | 17Networks_LH_LimbicA_TempPole_2 | -0.218119405 | -0.019624515 |
| 116 | 17Networks_LH_LimbicA_TempPole_3 | -0.072889282 | |
| 117 | 17Networks_LH_LimbicA_TempPole_4 | -0.049655003 | 0.003172451 |
| 118 | 17Networks_LH_LimbicA_TempPole_5 | 0.050483532 | |
| 119 | 17Networks_LH_LimbicA_TempPole_6 | -0.063024291 | |
| 120 | 17Networks_LH_LimbicA_TempPole_7 | -0.049971418 | |
| 121 | 17Networks_LH_ContA_Temp_1 | -0.172835721 | -0.077661268 |
| 122 | 17Networks_LH_ContA_IPS_1 | -0.121407713 | -0.000870215 |
| 123 | 17Networks_LH_ContA_IPS_2 | 0.057680714 | |
| 124 | 17Networks_LH_ContA_IPS_3 | 0.012420901 | |
| 125 | 17Networks_LH_ContA_IPS_4 | 0.047000433 | 0.022519872 |
| 126 | 17Networks_LH_ContA_IPS_5 | 0.167104363 | |
| 127 | 17Networks_LH_ContA_PFCd_1 | -0.169374106 | |
| 128 | 17Networks_LH_ContA_PFClv_1 | 0.091361331 | 0.026493277 |
| 129 | 17Networks_LH_ContA_PFClv_2 | 0.053773144 | |

| | | | |
|---|---|---|---|
| 130 | 17Networks_LH_ContA_PFCl_1 | 0.244231695 | 0.012930135 |
| 131 | 17Networks_LH_ContA_PFCl_2 | -0.132566316 | |
| 132 | 17Networks_LH_ContA_PFCl_3 | 0.200698596 | |
| 133 | 17Networks_LH_ContA_Cingm_1 | -0.011544625 | |
| 134 | 17Networks_LH_ContB_Temp_1 | 0.035498855 | |
| 135 | 17Networks_LH_ContB_Temp_2 | 0.144640302 | -0.021366656 |
| 136 | 17Networks_LH_ContB_IPL_1 | 0.07519101 | |
| 137 | 17Networks_LH_ContB_IPL_2 | -0.092744923 | |
| 138 | 17Networks_LH_ContB_IPL_3 | 0.046857667 | |
| 139 | 17Networks_LH_ContB_PFCd_1 | -0.178990291 | -0.02176958 |
| 140 | 17Networks_LH_ContB_PFClv_1 | 0.056984396 | -0.006052899 |
| 141 | 17Networks_LH_ContB_PFClv_2 | 0.098775975 | |
| 142 | 17Networks_LH_ContB_PFClv_3 | -0.089665704 | -0.010692169 |
| 143 | 17Networks_LH_ContB_PFCmp_1 | 0.210205983 | |
| 144 | 17Networks_LH_ContC_pCun_1 | 0.077631117 | |
| 145 | 17Networks_LH_ContC_pCun_2 | -0.032681411 | |
| 146 | 17Networks_LH_ContC_pCun_3 | -0.068349752 | |
| 147 | 17Networks_LH_ContC_Cingp_1 | 0.160601713 | |
| 148 | 17Networks_LH_ContC_Cingp_2 | -0.044163693 | |
| 149 | 17Networks_LH_DefaultA_IPL_1 | 0.011956638 | |
| 150 | 17Networks_LH_DefaultA_IPL_2 | 0.162119537 | |
| 151 | 17Networks_LH_DefaultA_PFCd_1 | -0.026018512 | |
| 152 | 17Networks_LH_DefaultA_PFCd_2 | 0.11103412 | |
| 153 | 17Networks_LH_DefaultA_PFCd_3 | 0.080145441 | |
| 154 | 17Networks_LH_DefaultA_pCunPCC_1 | 0.11064542 | |
| 155 | 17Networks_LH_DefaultA_pCunPCC_2 | -0.180814534 | |
| 156 | 17Networks_LH_DefaultA_pCunPCC_3 | 0.16252694 | |
| 157 | 17Networks_LH_DefaultA_pCunPCC_4 | 0.130635755 | |
| 158 | 17Networks_LH_DefaultA_pCunPCC_5 | 0.019312525 | 0.041483306 |
| 159 | 17Networks_LH_DefaultA_pCunPCC_6 | 0.056914491 | |
| 160 | 17Networks_LH_DefaultA_pCunPCC_7 | -0.105980108 | -0.07815911 |
| 161 | 17Networks_LH_DefaultA_PFCm_1 | -0.25484909 | |
| 162 | 17Networks_LH_DefaultA_PFCm_2 | -0.064099078 | |
| 163 | 17Networks_LH_DefaultA_PFCm_3 | 0.031761178 | |
| 164 | 17Networks_LH_DefaultA_PFCm_4 | 0.015158791 | 0.051894458 |
| 165 | 17Networks_LH_DefaultA_PFCm_5 | -0.055863846 | -0.01603468 |
| 166 | 17Networks_LH_DefaultA_PFCm_6 | 0.047837251 | -0.02237069 |
| 167 | 17Networks_LH_DefaultB_Temp_1 | 0.032041677 | 0.041235528 |
| 168 | 17Networks_LH_DefaultB_Temp_2 | 0.070461342 | -0.000496639 |
| 169 | 17Networks_LH_DefaultB_Temp_3 | -0.222924015 | -0.05147831 |
| 170 | 17Networks_LH_DefaultB_Temp_4 | 0.062524715 | |
| 171 | 17Networks_LH_DefaultB_Temp_5 | 0.001172208 | |
| 172 | 17Networks_LH_DefaultB_Temp_6 | 0.027804323 | 0.024112214 |
| 173 | 17Networks_LH_DefaultB_IPL_1 | -0.052049936 | 0.001735606 |

| | | | |
|---|---|---|---|
| 174 | 17Networks_LH_DefaultB_IPL_2 | 0.056094405 | |
| 175 | 17Networks_LH_DefaultB_PFCd_1 | -0.003741987 | |
| 176 | 17Networks_LH_DefaultB_PFCd_2 | 0.118346546 | 0.080636329 |
| 177 | 17Networks_LH_DefaultB_PFCd_3 | 0.172941138 | |
| 178 | 17Networks_LH_DefaultB_PFCd_4 | -0.11618996 | |
| 179 | 17Networks_LH_DefaultB_PFCd_5 | -0.114116058 | |
| 180 | 17Networks_LH_DefaultB_PFCd_6 | 0.000735951 | -0.000215671 |
| 181 | 17Networks_LH_DefaultB_PFCl_1 | -0.239298227 | -0.010949791 |
| 182 | 17Networks_LH_DefaultB_PFCl_2 | -0.16809535 | |
| 183 | 17Networks_LH_DefaultB_PFCv_1 | 0.028450399 | -0.086511435 |
| 184 | 17Networks_LH_DefaultB_PFCv_2 | 0.016459821 | |
| 185 | 17Networks_LH_DefaultB_PFCv_3 | 0.148960791 | -0.017596606 |
| 186 | 17Networks_LH_DefaultB_PFCv_4 | 0.053321994 | |
| 187 | 17Networks_LH_DefaultB_PFCv_5 | -0.013861087 | |
| 188 | 17Networks_LH_DefaultC_IPL_1 | -0.042943867 | 0.028749244 |
| 189 | 17Networks_LH_DefaultC_Rsp_1 | -0.084295715 | |
| 190 | 17Networks_LH_DefaultC_Rsp_2 | -0.100936657 | 0.006143675 |
| 191 | 17Networks_LH_DefaultC_Rsp_3 | -0.052271515 | |
| 192 | 17Networks_LH_DefaultC_PHC_1 | 0.18875291 | |
| 193 | 17Networks_LH_DefaultC_PHC_2 | 0.208105284 | |
| 194 | 17Networks_LH_DefaultC_PHC_3 | 0.217613884 | |
| 195 | 17Networks_LH_TempPar_1 | 0.011117901 | |
| 196 | 17Networks_LH_TempPar_2 | -0.110113974 | |
| 197 | 17Networks_LH_TempPar_3 | -0.008817069 | 0.027350809 |
| 198 | 17Networks_LH_TempPar_4 | 0.030899035 | 0.055184054 |
| 199 | 17Networks_LH_TempPar_5 | -0.064377733 | |
| 200 | 17Networks_LH_TempPar_6 | 0.016928516 | |
| 201 | 17Networks_RH_VisCent_ExStr_1 | 0.107169796 | 0.041759861 |
| 202 | 17Networks_RH_VisCent_ExStr_2 | 0.100178471 | |
| 203 | 17Networks_RH_VisCent_ExStr_3 | -0.247362998 | |
| 204 | 17Networks_RH_VisCent_ExStr_4 | 0.334497158 | |
| 205 | 17Networks_RH_VisCent_ExStr_5 | 0.289753436 | |
| 206 | 17Networks_RH_VisCent_ExStr_6 | -0.108678869 | -0.005802085 |
| 207 | 17Networks_RH_VisCent_Striate_1 | -0.042242145 | |
| 208 | 17Networks_RH_VisCent_ExStr_7 | -0.248833104 | |
| 209 | 17Networks_RH_VisCent_ExStr_8 | -0.088968907 | -0.076729822 |
| 210 | 17Networks_RH_VisCent_ExStr_9 | -0.060963787 | |
| 211 | 17Networks_RH_VisCent_ExStr_10 | -0.023121899 | 0.116829946 |
| 212 | 17Networks_RH_VisCent_ExStr_11 | -0.154653912 | |
| 213 | 17Networks_RH_VisPeri_ExStrInf_1 | 0.132820924 | -0.004366793 |
| 214 | 17Networks_RH_VisPeri_ExStrInf_2 | 0.227300636 | |
| 215 | 17Networks_RH_VisPeri_ExStrInf_3 | -0.052326009 | |
| 216 | 17Networks_RH_VisPeri_ExStrInf_4 | 0.103874378 | |
| 217 | 17Networks_RH_VisPeri_ExStrInf_5 | 0.021892981 | -0.040497591 |

| | | | |
|---|---|---|---|
| 218 | 17Networks_RH_VisPeri_StriCal_1 | 0.071474745 | |
| 219 | 17Networks_RH_VisPeri_StriCal_2 | -0.166912843 | -0.034693421 |
| 220 | 17Networks_RH_VisPeri_ExStrSup_1 | -0.224679345 | |
| 221 | 17Networks_RH_VisPeri_ExStrSup_2 | 0.160726471 | |
| 222 | 17Networks_RH_VisPeri_ExStrSup_3 | 0.01431292 | |
| 223 | 17Networks_RH_VisPeri_ExStrSup_4 | -0.014465924 | |
| 224 | 17Networks_RH_SomMotA_1 | -0.070137803 | |
| 225 | 17Networks_RH_SomMotA_2 | -0.256502043 | |
| 226 | 17Networks_RH_SomMotA_3 | 0.099499533 | |
| 227 | 17Networks_RH_SomMotA_4 | -0.084112494 | |
| 228 | 17Networks_RH_SomMotA_5 | 0.086928062 | |
| 229 | 17Networks_RH_SomMotA_6 | -0.177262562 | 0.004363984 |
| 230 | 17Networks_RH_SomMotA_7 | 0.000120315 | |
| 231 | 17Networks_RH_SomMotA_8 | -0.224858973 | -0.013012682 |
| 232 | 17Networks_RH_SomMotA_9 | -0.107665707 | |
| 233 | 17Networks_RH_SomMotA_10 | 0.205881304 | 0.019809304 |
| 234 | 17Networks_RH_SomMotA_11 | 0.126726999 | |
| 235 | 17Networks_RH_SomMotA_12 | 0.094335787 | |
| 236 | 17Networks_RH_SomMotA_13 | -0.119149695 | -0.046282367 |
| 237 | 17Networks_RH_SomMotA_14 | -0.144717013 | |
| 238 | 17Networks_RH_SomMotA_15 | 0.129692664 | |
| 239 | 17Networks_RH_SomMotA_16 | 0.169137041 | 0.086124243 |
| 240 | 17Networks_RH_SomMotA_17 | -0.027804884 | 0.047625373 |
| 241 | 17Networks_RH_SomMotA_18 | 0.085062539 | 0.013992048 |
| 242 | 17Networks_RH_SomMotA_19 | -0.098698034 | 0.01883836 |
| 243 | 17Networks_RH_SomMotA_20 | -0.20773655 | -0.001642863 |
| 244 | 17Networks_RH_SomMotB_Aud_1 | 0.108175875 | -0.000795472 |
| 245 | 17Networks_RH_SomMotB_Aud_2 | -0.187195915 | -0.003219342 |
| 246 | 17Networks_RH_SomMotB_Ins_1 | 0.115659323 | |
| 247 | 17Networks_RH_SomMotB_S2_1 | -0.099335349 | |
| 248 | 17Networks_RH_SomMotB_S2_2 | -0.012072391 | 0.010492558 |
| 249 | 17Networks_RH_SomMotB_Aud_3 | 0.060878484 | -0.011328966 |
| 250 | 17Networks_RH_SomMotB_S2_3 | 0.125278117 | |
| 251 | 17Networks_RH_SomMotB_S2_4 | -0.053410532 | |
| 252 | 17Networks_RH_SomMotB_S2_5 | -0.135228791 | |
| 253 | 17Networks_RH_SomMotB_S2_6 | 0.098391472 | 0.012091215 |
| 254 | 17Networks_RH_SomMotB_S2_7 | -0.051736647 | |
| 255 | 17Networks_RH_SomMotB_S2_8 | -0.108249385 | 0.033522499 |
| 256 | 17Networks_RH_SomMotB_Cent_1 | 0.047559601 | 0.002127041 |
| 257 | 17Networks_RH_SomMotB_Cent_2 | 0.075405352 | |
| 258 | 17Networks_RH_SomMotB_Cent_3 | -0.044475135 | |
| 259 | 17Networks_RH_DorsAttnA_TempOcc_1 | 0.011011557 | |
| 260 | 17Networks_RH_DorsAttnA_TempOcc_2 | -0.018168437 | |
| 261 | 17Networks_RH_DorsAttnA_TempOcc_3 | -0.12782069 | 0.001917617 |

| | | | |
|---|---|---|---|
| 262 | 17Networks_RH_DorsAttnA_ParOcc_1 | 0.124002667 | |
| 263 | 17Networks_RH_DorsAttnA_ParOcc_2 | 0.153096145 | |
| 264 | 17Networks_RH_DorsAttnA_ParOcc_3 | 0.051774567 | |
| 265 | 17Networks_RH_DorsAttnA_SPL_1 | -0.027765677 | |
| 266 | 17Networks_RH_DorsAttnA_SPL_2 | -0.082243294 | |
| 267 | 17Networks_RH_DorsAttnA_SPL_3 | -0.028277134 | |
| 268 | 17Networks_RH_DorsAttnA_SPL_4 | 0.133975723 | |
| 269 | 17Networks_RH_DorsAttnA_SPL_5 | 0.025486856 | -0.023832507 |
| 270 | 17Networks_RH_DorsAttnA_SPL_6 | -0.222978842 | 0.031216895 |
| 271 | 17Networks_RH_DorsAttnA_SPL_7 | 0.332533633 | |
| 272 | 17Networks_RH_DorsAttnA_SPL_8 | -0.012939089 | -0.039753931 |
| 273 | 17Networks_RH_DorsAttnB_TempOcc_1 | 0.128015541 | |
| 274 | 17Networks_RH_DorsAttnB_PostC_1 | 0.137554253 | |
| 275 | 17Networks_RH_DorsAttnB_PostC_2 | 0.098282929 | -0.021131876 |
| 276 | 17Networks_RH_DorsAttnB_PostC_3 | -0.146697627 | |
| 277 | 17Networks_RH_DorsAttnB_PostC_4 | -0.149766241 | |
| 278 | 17Networks_RH_DorsAttnB_PostC_5 | 0.153414057 | -0.014971212 |
| 279 | 17Networks_RH_DorsAttnB_PostC_6 | 0.165017216 | -0.035829591 |
| 280 | 17Networks_RH_DorsAttnB_PostC_7 | 0.170005663 | |
| 281 | 17Networks_RH_DorsAttnB_PostC_8 | -0.045181215 | -0.000640858 |
| 282 | 17Networks_RH_DorsAttnB_FEF_1 | -0.195685225 | 0.053740179 |
| 283 | 17Networks_RH_DorsAttnB_FEF_2 | 0.106014939 | |
| 284 | 17Networks_RH_DorsAttnB_FEF_3 | -0.218198265 | |
| 285 | 17Networks_RH_SalVentAttnA_ParOper_1 | 0.166416738 | |
| 286 | 17Networks_RH_SalVentAttnA_ParOper_2 | -0.206800881 | |
| 287 | 17Networks_RH_SalVentAttnA_ParOper_3 | 0.080210511 | -0.004177292 |
| 288 | 17Networks_RH_SalVentAttnA_PrC_1 | 0.029420196 | -0.000955279 |
| 289 | 17Networks_RH_SalVentAttnA_Ins_1 | 0.025061896 | 0.061357847 |
| 290 | 17Networks_RH_SalVentAttnA_Ins_2 | -0.034573136 | -0.020927285 |
| 291 | 17Networks_RH_SalVentAttnA_Ins_3 | -0.023098664 | |
| 292 | 17Networks_RH_SalVentAttnA_Ins_4 | -0.091253875 | |
| 293 | 17Networks_RH_SalVentAttnA_FrOper_1 | -0.165125838 | -0.086221628 |
| 294 | 17Networks_RH_SalVentAttnA_FrOper_2 | 0.097997741 | |
| 295 | 17Networks_RH_SalVentAttnA_FrOper_3 | -0.073186254 | |
| 296 | 17Networks_RH_SalVentAttnA_FrMed_1 | 0.046197583 | |
| 297 | 17Networks_RH_SalVentAttnA_ParMed_1 | -0.201289461 | |
| 298 | 17Networks_RH_SalVentAttnA_ParMed_2 | 0.095985368 | 0.024554491 |
| 299 | 17Networks_RH_SalVentAttnA_FrMed_2 | 0.175759132 | |
| 300 | 17Networks_RH_SalVentAttnA_ParMed_3 | -0.017246542 | |
| 301 | 17Networks_RH_SalVentAttnA_ParMed_4 | 0.049498929 | -0.004997533 |
| 302 | 17Networks_RH_SalVentAttnA_FrMed_3 | -0.201151913 | |
| 303 | 17Networks_RH_SalVentAttnA_FrMed_4 | 0.009566472 | |
| 304 | 17Networks_RH_SalVentAttnB_IPL_1 | 0.189518399 | |
| 305 | 17Networks_RH_SalVentAttnB_PFClv_1 | -0.091437054 | |

| | | | |
|---|---|---|---|
| 306 | 17Networks_RH_SalVentAttnB_PFCl_1 | 0.183471736 | 0.001664832 |
| 307 | 17Networks_RH_SalVentAttnB_PFCl_2 | 0.123044795 | -0.024237492 |
| 308 | 17Networks_RH_SalVentAttnB_PFCl_3 | -0.109553782 | 0.000210713 |
| 309 | 17Networks_RH_SalVentAttnB_Ins_1 | -0.199573606 | |
| 310 | 17Networks_RH_SalVentAttnB_Ins_2 | 0.079729965 | |
| 311 | 17Networks_RH_SalVentAttnB_PFCmp_1 | -0.024876011 | |
| 312 | 17Networks_RH_SalVentAttnB_PFCmp_2 | 0.091453831 | 0.004392898 |
| 313 | 17Networks_RH_LimbicB_OFC_1 | -0.052568068 | |
| 314 | 17Networks_RH_LimbicB_OFC_2 | -0.055055836 | |
| 315 | 17Networks_RH_LimbicB_OFC_3 | 0.113416662 | |
| 316 | 17Networks_RH_LimbicB_OFC_4 | -0.029259337 | |
| 317 | 17Networks_RH_LimbicB_OFC_5 | -0.008505607 | |
| 318 | 17Networks_RH_LimbicB_OFC_6 | 0.255407605 | |
| 319 | 17Networks_RH_LimbicA_TempPole_1 | -0.078842426 | |
| 320 | 17Networks_RH_LimbicA_TempPole_2 | 0.050843044 | |
| 321 | 17Networks_RH_LimbicA_TempPole_3 | -0.049271485 | |
| 322 | 17Networks_RH_LimbicA_TempPole_4 | 0.170302679 | |
| 323 | 17Networks_RH_LimbicA_TempPole_5 | -0.069862031 | |
| 324 | 17Networks_RH_LimbicA_TempPole_6 | 0.260749002 | -0.027530927 |
| 325 | 17Networks_RH_ContA_IPS_1 | -0.192432639 | -0.017015243 |
| 326 | 17Networks_RH_ContA_IPS_2 | -0.042779453 | |
| 327 | 17Networks_RH_ContA_IPS_3 | -0.02591333 | |
| 328 | 17Networks_RH_ContA_IPS_4 | 0.005553417 | 0.003028594 |
| 329 | 17Networks_RH_ContA_PFCd_1 | -0.117592253 | -0.053974212 |
| 330 | 17Networks_RH_ContA_PFCl_1 | 0.025832819 | |
| 331 | 17Networks_RH_ContA_PFCl_2 | -0.230795204 | |
| 332 | 17Networks_RH_ContA_PFCl_3 | 0.153906284 | |
| 333 | 17Networks_RH_ContA_PFCl_4 | -0.023516128 | -0.000465008 |
| 334 | 17Networks_RH_ContA_PFCl_5 | 0.032323607 | 0.037568721 |
| 335 | 17Networks_RH_ContA_Cingm_1 | 0.001664471 | |
| 336 | 17Networks_RH_ContB_Temp_1 | 0.022749357 | |
| 337 | 17Networks_RH_ContB_Temp_2 | -0.125646254 | |
| 338 | 17Networks_RH_ContB_IPL_1 | -0.065902756 | |
| 339 | 17Networks_RH_ContB_IPL_2 | -0.08128531 | -0.042026615 |
| 340 | 17Networks_RH_ContB_IPL_3 | -0.098219473 | |
| 341 | 17Networks_RH_ContB_IPL_4 | -0.021763252 | |
| 342 | 17Networks_RH_ContB_PFCld_1 | -0.204332797 | |
| 343 | 17Networks_RH_ContB_PFCld_2 | -0.049098014 | |
| 344 | 17Networks_RH_ContB_PFCld_3 | -0.159463909 | 0.051075803 |
| 345 | 17Networks_RH_ContB_PFCld_4 | -0.102860698 | -0.033115018 |
| 346 | 17Networks_RH_ContB_PFClv_1 | 0.023791256 | 0.035704684 |
| 347 | 17Networks_RH_ContB_PFClv_2 | -0.166869781 | -0.071580959 |
| 348 | 17Networks_RH_ContB_PFClv_3 | -0.015363162 | 0.043272934 |
| 349 | 17Networks_RH_ContB_PFClv_4 | 0.245744425 | |

| | | | |
|---|---|---|---|
| 350 | 17Networks_RH_ContB_PFCmp_1 | 0.096958716 | |
| 351 | 17Networks_RH_ContC_pCun_1 | -0.113899698 | 0.05449326 |
| 352 | 17Networks_RH_ContC_pCun_2 | 0.075885945 | |
| 353 | 17Networks_RH_ContC_pCun_3 | -0.179432413 | |
| 354 | 17Networks_RH_ContC_pCun_4 | 0.034313096 | |
| 355 | 17Networks_RH_ContC_pCun_5 | -0.284415046 | |
| 356 | 17Networks_RH_ContC_Cingp_1 | -0.049772768 | |
| 357 | 17Networks_RH_ContC_Cingp_2 | -0.235340341 | |
| 358 | 17Networks_RH_DefaultA_Temp_1 | -0.006506102 | |
| 359 | 17Networks_RH_DefaultA_IPL_1 | 0.014961579 | 0.004584649 |
| 360 | 17Networks_RH_DefaultA_IPL_2 | 0.145230251 | -0.047293527 |
| 361 | 17Networks_RH_DefaultA_PFCd_1 | -0.071397833 | 0.022725311 |
| 362 | 17Networks_RH_DefaultA_PFCd_2 | 0.037317798 | |
| 363 | 17Networks_RH_DefaultA_pCunPCC_1 | -0.011695142 | |
| 364 | 17Networks_RH_DefaultA_pCunPCC_2 | -0.060356443 | |
| 365 | 17Networks_RH_DefaultA_pCunPCC_3 | -0.016964468 | -0.013743166 |
| 366 | 17Networks_RH_DefaultA_pCunPCC_4 | -0.04453629 | 0.037575132 |
| 367 | 17Networks_RH_DefaultA_pCunPCC_5 | 0.199671108 | |
| 368 | 17Networks_RH_DefaultA_PFCm_1 | -0.021135988 | |
| 369 | 17Networks_RH_DefaultA_PFCm_2 | 0.109718081 | |
| 370 | 17Networks_RH_DefaultA_PFCm_3 | -0.158746364 | -0.004338126 |
| 371 | 17Networks_RH_DefaultA_PFCm_4 | -0.095962307 | |
| 372 | 17Networks_RH_DefaultA_PFCm_5 | -0.020260655 | |
| 373 | 17Networks_RH_DefaultA_PFCm_6 | -0.188371343 | |
| 374 | 17Networks_RH_DefaultB_Temp_1 | -0.030367076 | |
| 375 | 17Networks_RH_DefaultB_Temp_2 | 0.207460591 | |
| 376 | 17Networks_RH_DefaultB_AntTemp_1 | 0.066680729 | |
| 377 | 17Networks_RH_DefaultB_PFCd_1 | 0.099397038 | 0.008143918 |
| 378 | 17Networks_RH_DefaultB_PFCd_2 | 0.226751887 | |
| 379 | 17Networks_RH_DefaultB_PFCd_3 | -0.034181797 | |
| 380 | 17Networks_RH_DefaultB_PFCd_4 | 0.045213037 | 0.024142518 |
| 381 | 17Networks_RH_DefaultB_PFCd_5 | -0.048424626 | |
| 382 | 17Networks_RH_DefaultB_PFCv_1 | -0.065869401 | |
| 383 | 17Networks_RH_DefaultB_PFCv_2 | 0.045630326 | 0.019491153 |
| 384 | 17Networks_RH_DefaultB_PFCv_3 | 0.107117464 | |
| 385 | 17Networks_RH_DefaultC_IPL_1 | -0.106828581 | |
| 386 | 17Networks_RH_DefaultC_IPL_2 | -0.167035084 | |
| 387 | 17Networks_RH_DefaultC_Rsp_1 | 0.237601492 | |
| 388 | 17Networks_RH_DefaultC_Rsp_2 | 0.168654649 | |
| 389 | 17Networks_RH_DefaultC_PHC_1 | -0.000242299 | |
| 390 | 17Networks_RH_DefaultC_PHC_2 | 0.026772439 | |
| 391 | 17Networks_RH_TempPar_1 | 0.134868531 | |
| 392 | 17Networks_RH_TempPar_2 | -0.078043608 | |
| 393 | 17Networks_RH_TempPar_3 | 0.136224059 | |

| | | | |
|---|---|---|---|
| 394 | 17Networks_RH_TempPar_4 | -0.109410462 | |
| 395 | 17Networks_RH_TempPar_5 | 0.103258275 | |
| 396 | 17Networks_RH_TempPar_6 | 0.023967572 | |
| 397 | 17Networks_RH_TempPar_7 | -0.215875491 | |
| 398 | 17Networks_RH_TempPar_8 | -0.103727092 | |
| 399 | 17Networks_RH_TempPar_9 | 0.077884963 | |
| 400 | 17Networks_RH_TempPar_10 | 0.13031885 | -0.010321144 |

*Note.* LH = left hemisphere, RH = right hemisphere, VisCent = central visual, VisPeri = peripheral visual, SomMot = somatomotor, DorsAttn = dorsal attention, SalVentAttn = salience/ventral attention, Cont = frontoparietal control, TempPar = temporoparietal, *g* = general intelligence, O/I = Openness/Intellect

**Table A2**

*Performance metrics of models using flexibility of all parcels as predictors in the γ = 1 partition*

| Criterion | Type | Training Sample | | | | | Test Sample | | |
|---|---|---|---|---|---|---|---|---|---|
| | | α | λ | RMSE | MAE | $R^2$ | RMSE | MAE | $R^2$ |
| O/I | Overall | 0.1 | .0090283 | 1.87 | 1.48 | .040 | 1.84 | 1.45 | .001 |
| | Cohesive | 1 | .1101182 | 1.01 | .79 | .032 | .99 | .80 | .000 |
| | Disjoint | 0.8 | .1575058 | .99 | .78 | .066 | 1.00 | .80 | .000 |
| *g* | Overall | 0.3 | .0000587 | 3.12 | 2.45 | .023 | 2.40 | 1.91 | .003 |
| | Cohesive | 1 | .0518615 | 1.05 | .83 | .025 | 1.03 | .84 | .001 |
| | Disjoint | 0.9 | .0579943 | 1.02 | .80 | .020 | 1.02 | .83 | .001 |

*Note.* *g* = general intelligence. O/I = Openness/Intellect. α = mixing parameter, λ = penalty parameter, RMSE = root-mean-square error. MAE = mean absolute error

### Predicting Plasticity from Parcel Flexibility

Metatrait Plasticity scores were computed by averaging participants' Extraversion and Openness/Intellect scores. Scores of the metatrait Stability were also computed by averaging Conscientiousness, Agreeableness, and the inverse of Neuroticism scores to be included as a covariate. We repeated the procedure described in the main text predicting Plasticity scores from each of the flexibility measures. These models included age, gender, mean relative RMS movement, handedness, total brain volume, the type of image reconstruction algorithm, *g,* and metatrait Stability scores as covariates. Stability was included as a covariate to account for the influence of evaluative consistency bias (Feeley 2002; Anusic et al. 2009). Results from these tests are described in Table A3.

**Table A3**

*Performance metrics of models predicting metatrait Plasticity from parcel flexibility*

| Community | Type | Training Sample | | | | | Test Sample | | |
|---|---|---|---|---|---|---|---|---|---|
| | | α | λ | RMSE | MAE | $R^2$ | RMSE | MAE | $R^2$ |
| γ = 1.1 | Overall | 1 | .1079578 | 1.00 | .78 | .032 | 1.00 | .78 | .002 |
| | Cohesive | 0.8 | .0967937 | 1.01 | .79 | .050 | 1.00 | .79 | .000 |
| | Disjoint | 1 | .0214803 | 1.17 | .93 | .029 | 1.10 | .88 | .004 |
| γ = 1.2 | Overall | 0.1 | .1132511 | 1.08 | .84 | .040 | 1.17 | .93 | .000 |
| | Cohesive | 0.7 | .0000599 | 2.48 | 2.00 | .041 | 2.89 | 2.25 | .001 |
| | Disjoint | 0.1 | .1176264 | 1.15 | .91 | .019 | 1.17 | .93 | .000 |

*Note.* α = mixing parameter, λ = penalty parameter, RMSE = root-mean-square error. MAE = mean absolute error
